# Negative allosteric modulation of the µ-opioid receptor

**DOI:** 10.1101/2023.09.08.556921

**Authors:** Evan S. O’Brien, Vipin Ashok Rangari, Amal El Daibani, Shainnel O. Eans, Betsy White, Haoqing Wang, Yuki Shiimura, Kaavya Krishna Kumar, Kevin Appourchaux, Weijiao Huang, Chensong Zhang, Jesper M. Mathiesen, Tao Che, Jay P. McLaughlin, Susruta Majumdar, Brian K. Kobilka

**Affiliations:** Department of Molecular and Cellular Physiology, Stanford University School of Medicine, 279 Campus Drive, Stanford, CA 94305, USA; Center for Clinical Pharmacology, University of Health Sciences & Pharmacy at St Louis and Washington University School of Medicine, St. Louis, MO 63110, USA; Department of Pharmacodynamics, University of Florida, Gainesville, FL 32610, USA; Division of Molecular Genetics, Institute of Life Science, Kurume University, Fukuoka, Japan; Division of CryoEM and Bioimaging, SSRL, SLAC National Acceleration Laboratory, Menlo Park, CA 94025, USA; Department of Drug Design and Pharmacology, University of Copenhagen, Copenhagen 2100, Denmark

## Abstract

The µ-opioid receptor (µOR) is a well-established target for analgesia, yet conventional opioid receptor agonists cause serious adverse effects, notably addiction and respiratory depression, which have led to the present opioid overdose epidemic. µOR negative allosteric modulators (NAMs) may serve as powerful tools in preventing opioid overdose deaths, but promising chemical scaffolds remain elusive. We screened a large DNA-encoded chemical library against inactive µOR, counter-screening with active, G-protein and agonist bound receptor to “steer” selections toward functional negative allosteric modulators. We discovered a NAM compound with high and selective enrichment to inactive µOR; the molecule potently blocks the activity of orthosteric agonists and enhances the affinity of the key opioid overdose reversal molecule, naloxone. It accomplishes this by binding to a site on the extracellular vestibule proximal to naloxone, stabilizing a unique inactive conformation of the extracellular portions of the second and seventh transmembrane helices. The NAM perturbs orthosteric ligand kinetics in therapeutically desirable ways and works cooperatively with low doses of naloxone *in vivo* to inhibit morphine-induced antinociception, respiratory depression and conditioned place preference while minimizing withdrawal behaviors. Our results provide detailed structural insights into the mechanism of a negative allosteric modulator for the µOR and demonstrate how it can be exploited *in vivo*.

## Introduction

Traditional opioid compounds extracted from *Papaver somniferum* including morphine have been used both recreationally and as potent pain relief molecules for millennia. Related semi-synthetic derivates such as oxycodone have been extensively prescribed clinically as analgesics. These compounds represent best-in-class treatments for acute pain management^1^ yet have also been commonly prescribed as long-term analgesic treatments, where their high potential for addiction and abuse, in conjunction with their severe respiratory depression effects, have fueled the current opioid overdose epidemic^1–3^. While ever more patients were becoming addicted to over-prescribed opioids, fully synthetic (and much more potent) opioid agonists such as fentanyl have been exploited as cheap additives to recreationally-used opioid mixtures^4,5^. Naloxone (Narcan) is the most common and effective treatment for opioid overdoses, however, larger and repeated doses are needed in response to overdoses from more potent and long-lasting opioids such as fentanyl^5,6^.

All these molecules share a similar mode of action as orthosteric agonists of the µ-opioid receptor (µOR), albeit with different affinities for the receptor and extents of intracellular signaling (efficacies). Agonists and antagonists of the µOR bind at an overlapping orthosteric site in the extracellular vestibule of the receptor and share a set of key interactions with endogenous opioid signaling peptides, namely the enkephalins, endorphins and endomorphins^7–9^. Given the limitations of the current slate of orthosteric opioid molecules, as well as the severity of the current opioid overdose epidemic, new classes of µOR-modulating compounds with distinct mechanisms of action are highly desirable and have emerged as priorities of the National Institute of Drug Abuse (NIDA)^10^.

Rather than further explore agonism at the traditional orthosteric site, modulation of receptor activity via binding of molecules at alternate sites on the receptor (allosteric sites) potentially provides a series of advantages. Unique signaling bias as well as functional selectivity *in vivo* can be achieved by extending orthosteric molecules into allosteric pockets^11^. Allosteric modulators may act more specifically through the µOR (rather than its κ-opioid receptor (κOR) and δ-opioid receptor (δOR) counterparts) due to decreased evolutionary similarity outside of the conserved opioid receptor orthosteric pocket. Allosteric modulators can sometimes display “probe dependence”, preferentially modulating the activity of one orthosteric molecule over others^12^. For instance, NAMs that are more effective against exogenous than endogenous agonists would be highly desirable. Certain small molecules, including cannabinoids^13^ and the selective κOR agonist salvinorin A^14^, have been identified as NAMs of the µOR, though their potencies for their selective targets (cannabinoid receptor 1 (CB1) and κOR) are much higher than those observed against the µOR (high µM), suggesting they are highly unlikely to display selective µOR-NAM activity *in vivo*. Selective, potent NAMs for the µOR have remained elusive.

We set out to identify such compounds through a directed screen of a large DNA-encoded chemical library (DEL) composed of a series of small fragments covalently linked together in a combinatorial manner (Fig. 1a) all conjugated to a DNA “barcode” sequence identifying the molecule to which it is linked. Due to this combinatorial nature, DELs can be composed of billions or even trillions of individual components covering large areas of chemical space^15–17^. This approach has been used to identify positive and negative allosteric modulators for the β_2_ adrenergic receptor^18–21^, though previous GPCR DEL selections have relied on a single receptor condition (e.g. agonist-bound receptor^18^), requiring synthesis of large numbers of compounds to identify those that bind target *and* have the desired biological activity. Our “steered” screening approach (Fig. 1a), centered around counter-screening against active, G_i_-bound µOR, resulted in the discovery of a potent µOR NAM from the synthesis of a single compound. Using cryo-electron microscopy (cryoEM), we demonstrate that the molecule “caps” the orthosteric antagonist naloxone in the orthosteric pocket and inhibits receptor activity by stabilizing a distinct inactive conformation of the extracellular vestibule. This activity translates to *in vivo* mouse models, where the NAM works cooperatively to boost the ability of low-doses of naloxone to reverse morphine induced effects while minimizing withdrawal symptoms.

**Figure 1.**
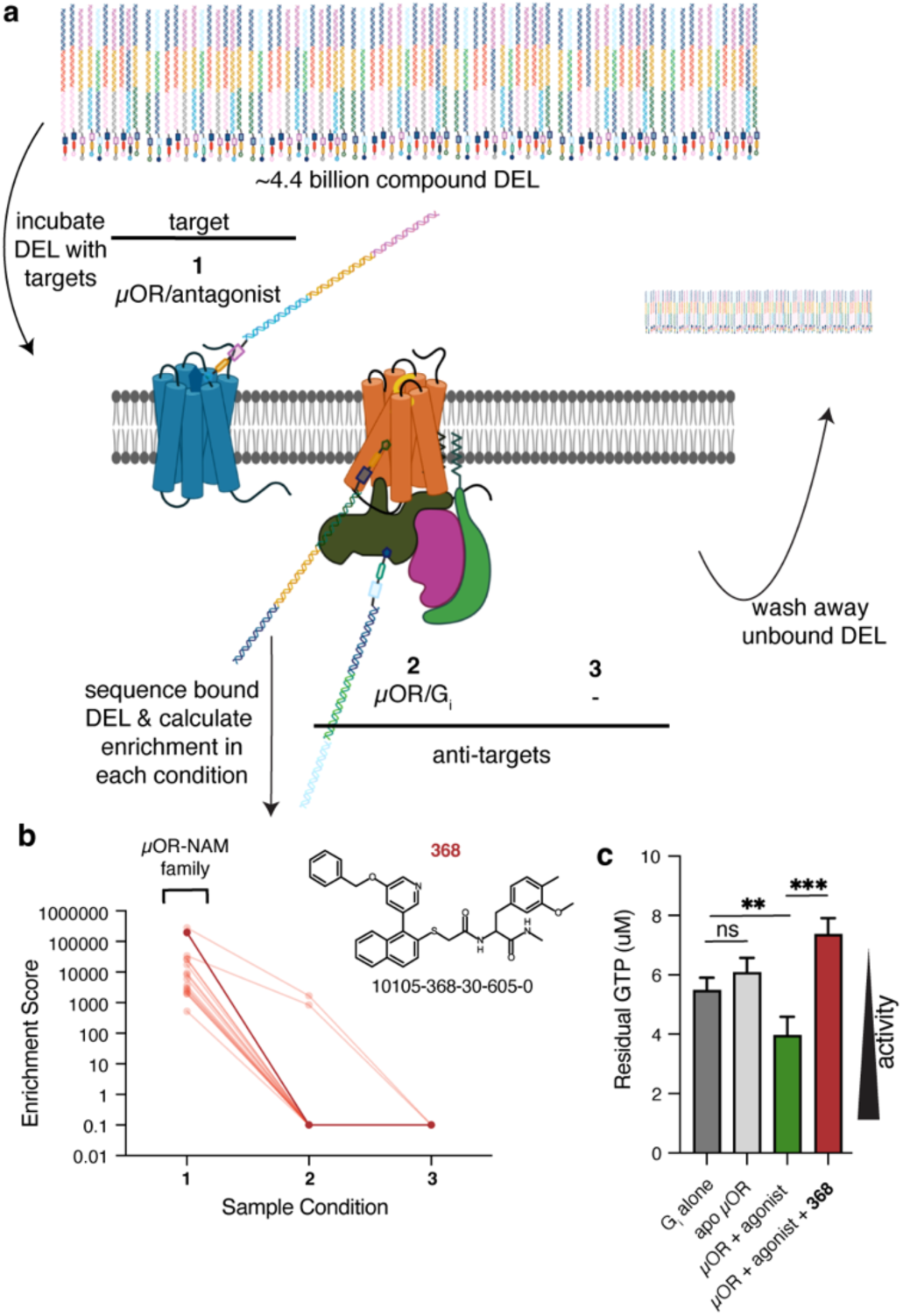
DEL screen for new µOR allosteric modulators. (**a**) Schematic detailing the DEL selection scheme including target (**1** - inactive µOR/naloxone), anti-target (**2** – active µOR/G_i_) and no target control (**3**). Figure created with BioRender. (**b**) Enrichment scores for **368** family members (semi-transparent) and chemical structure for the selected and characterized NAM (solid) across the 3 sample conditions. (**c**) GTP turnover assay used to initially assess the activity of the NAM. G_i_ alone has intrinsic GTPase activity (dark gray) that is not significantly impacted by the presence of apo µOR (light gray), but is enhanced when the receptor is bound to the full agonist peptide met-enkephalin (green). The NAM (red) significantly inhibits agonist-induced turnover. P values are denoted as follows: ns (P>0.05), * (P≤0.05), ** (P≤0.01), *** (P≤0.001), and **** (P≤0.0001) and were calculated using an unpaired t test assuming Gaussian distributions.

## Results

### Targeted DNA-encoded library screen for discovering GPCR NAMs

To discover NAMs that inhibit activation of the receptor, we selected for DEL components that bind specifically to µOR saturated with naloxone (condition **1**, Fig. 1a). To select against compounds that bind in a non-conformationally sensitive manner (silent allosteric modulators, SAMs) or that have some binding propensity to the active conformation of the receptor, we included µOR bound to agonist (met-enkephalin) and G_i_ as a fully-active anti-target (condition **2**, Fig. 1a). Finally, no target control (condition **3**, Fig. 1a) eliminates molecules that bind non-specifically to the beads used to immobilize receptors. Molecules that bind specifically to condition **1** and minimally to the others are likely to be µOR NAMs.

After several rounds of binding the DEL to all conditions and washing away unbound components, the enriched molecules were subjected to next-generation sequencing and relative enrichment scores were calculated based on the number of sequence reads in each condition (Fig. 1b). There were several “families” of enriched compounds in all conditions, an example of which is highlighted in Fig. 1b, with individual family members sharing one or two fragments in common (Table S1). All members are specifically enriched in the inactive form of µOR. We selected the best-enriched compound (10105-368-30-605; Compound **368**; calculated properties in Table S2) without detectable binding to anti-targets for further characterization. We first tested for activity using an *in vitro* GTP-turnover assay^22,23^ where we assessed the ability of **368** to dampen agonist-dependent GTP depletion. Indeed, **368** significantly inhibits met-enkephalin induced µOR activation of G_i_ (Fig. 1c). The observed inhibition of GTP turnover in the presence of NAM only occurs in the presence of receptor and G_i_ (Fig. S1a). Given the large (and conformationally selective) enrichment and observed activity of **368** as a µOR inhibitor, we were interested in more extensively characterizing its mechanism of action.

### Pharmacology of Compound 368

Using the GTP-depletion assay, we first showed that **368** inhibits µOR/met-enkephalin induced activation of G_i_ with an observed potency in the single-digit µM range (Fig. 2a), though the concentration of receptor is on the same scale, making quantitative evaluation of potencies difficult. The effect saturates at ∼10 µM **368**, nearly eliminating receptor induced GTP turnover (Fig. 2a). To assess the potency of **368** more quantitatively, we observed its effect on ^3^H-naloxone binding to µOR-expressing membranes. Consistent with its selective enrichment to inactive, naloxone-bound receptor and its inhibition of µOR induced inhibition of G_i_ signaling, titrating **368** results in increased binding of ^3^H-naloxone with an EC_50_ of 133 nM (Fig. 2b). This increase in binding is the result of a ∼2.6-fold increase in ^3^H-naloxone affinity in the presence of excess **368** (Fig. S1b). We next tested whether the observed biochemical cooperativity with naloxone was also present in cells. To that end, we conducted a TRUPATH^24^ assay in antagonist mode for µOR-expressing cells, titrating in naloxone to observe inhibition of G-protein heterotrimer dissociation using bioluminescence resonance energy transfer (BRET). The observed potency of naloxone for inhibiting agonist-induced signaling is enhanced by 7.6-fold in the presence of **368** (Fig. 2c, Table S3). This cooperativity with naloxone binding, observed biochemically and in cells, is also a direct demonstration of the allosteric mechanism of action of the compound, at least with respect to naloxone. **368** inhibits receptor-catalyzed GTP turnover in the absence of orthosteric ligand (Fig. 2d), designating the molecule as an inverse agonist NAM (inverse ago-NAM) for the µOR. It also inhibits additional turnover caused by the extremely weak partial agonist/neutral antagonist naloxone as well that from full agonist DAMGO (Fig. 2d). **368** results in inhibition of µOR-induced signaling when bound to a variety of orthosteric agonists (Fig. S1c), though excess concentrations of ligands are present.

**Figure 2.**
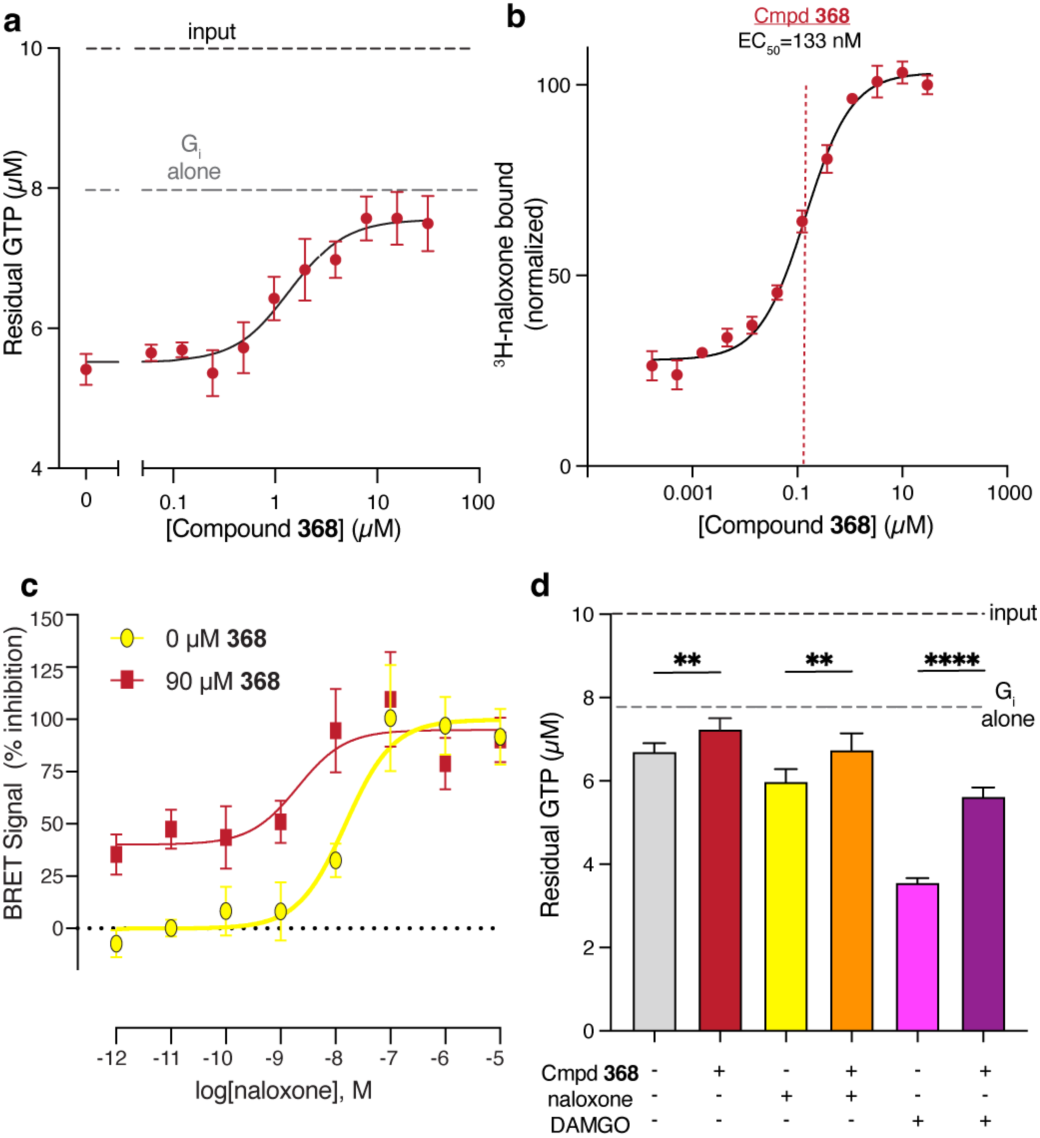
Cmpd 368 inhibits turnover cooperatively with naloxone. (**a**) Introduction of increasing amounts of **368** to the GTP turnover assay results in nearly full inhibition of excess (20 µM) met-enkephalin-bound receptor-mediated turnover with a potency closer to single-digit µM levels. (**b**) Increasing amounts of **368** result in an increase in antagonist (^3^H-naloxone) binding to the receptor with an observed EC_50_ of 133 nM (95% CI of 112 to 159 nM) (dashed red line), demonstrating the allosteric nature of the activity. (**c**) Reversal of G_i1_ activation by the µOR by titration of naloxone in the absence (yellow) or presence (red) of **368**. (**d**) GTP turnover assay with excess concentrations of **368**, antagonist naloxone, or full agonist DAMGO. **368** significantly inhibit basal signaling of the receptor (gray vs. red) as well as inhibiting both the extremely weak partial activity of naloxone (yellow vs. orange) or the full activity of DAMGO (pink vs. purple). P values are denoted as follows: ns (P>0.05), * (P≤0.05), ** (P≤0.01), *** (P≤0.001), and **** (P≤0.0001) and were calculated using an unpaired t test assuming Gaussian distributions.

We next conducted a series of in-cell TRUPATH^24^ assays to test the effect of **368** on G-protein activation in cells. We tested several orthosteric agonists (morphine, fentanyl, and met-enkephalin) and all Gα-protein family members through which µOR signals (G_i1_, G_i2_, G_i3_, G_oA_, G_oB_, and G_z_). The observed potency in the TRUPATH assays for all 3 ligands was decreased by more than 10-fold in the presence of **368** (Fig. 3a, Fig. 3b, Fig. 3c, Table S4) with all G_i/o_ subtypes inhibited in potency nearly equally (Fig. 3d) (i.e. there is no obvious biased negative allosterism of **368** across G_i_ family members). Agonist stimulation of receptors predominantly coupled to G_i/o_ family G-proteins results in decreased cAMP levels. We next tested whether the observed **368** inhibition of agonist potency for dissociating G-proteins translates to inhibition of agonist-induced decreases in cAMP levels. Indeed, titrating increasing concentrations of **368** at EC_80_ concentrations of orthosteric agonist results in a dose-dependent elevation of cAMP levels (Fig. 3e), albeit with lower NAM potencies than those observed biochemically (Fig. 2a, 2b).

**Figure 3.**
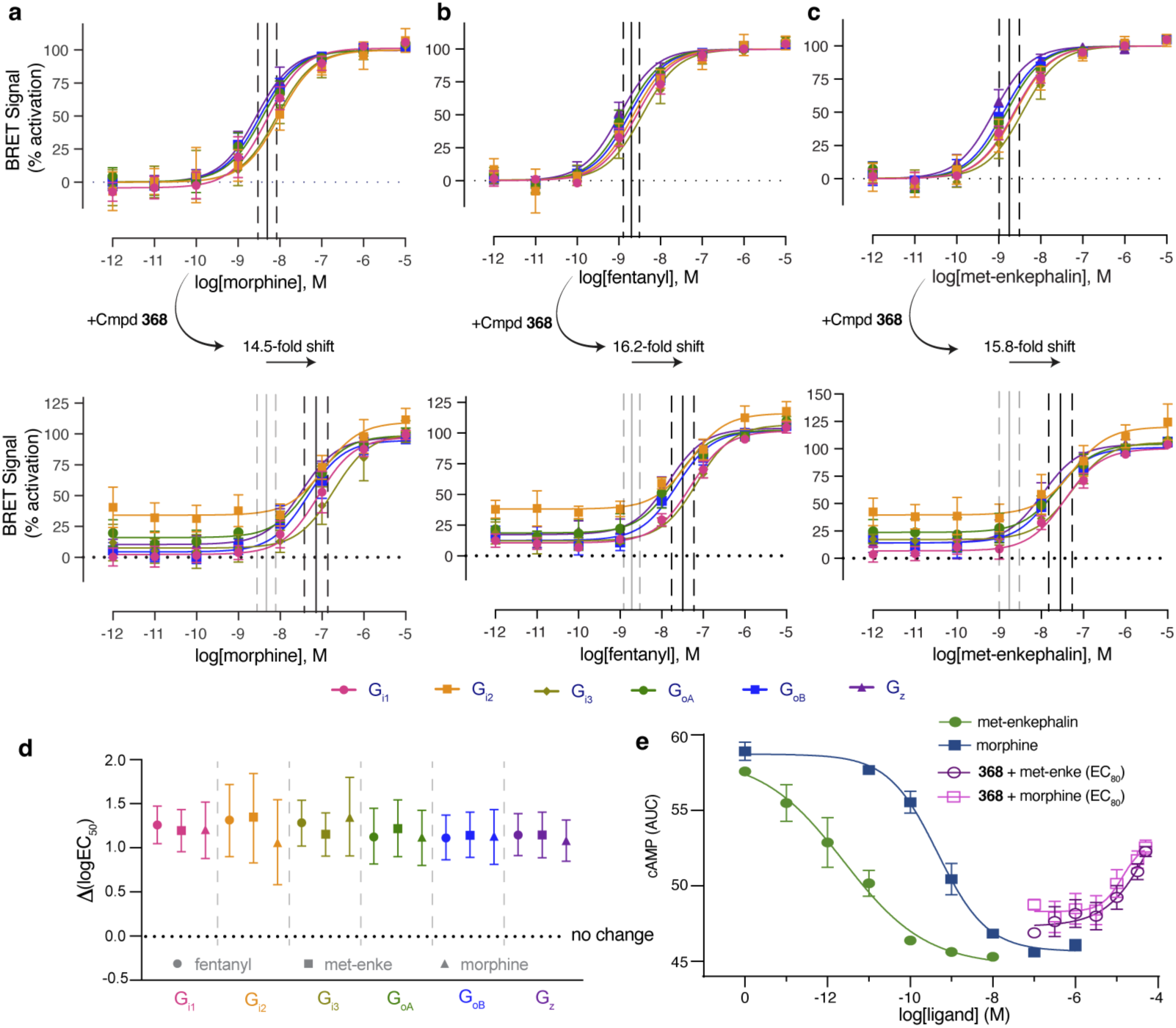
*In-cell* activity of 368. Titrations of (**a**) morphine, (**b**) fentanyl, and (**c**) met-enkephalin result in activation of an assortment of G-proteins (G_i1_, pink; G_i2_, orange; G_i3_, pale green; G_oA_, green; G_oB_, blue; and G_z_, purple) as observed in the TRUPATH assay by a change in BRET signal as the Gα and Gβγ subunits separate. Activation by all 3 agonists was also calculated in the presence of 90 µM **368** (bottom panels). The average log(EC_50_) for all 6 G-protein activation curves within each panel is shown as a black line (with dashed grey lines representing the standard deviation among different G protein subtypes). The average fold change in EC_50_ upon addition of **368** for morphine (14.5), fentanyl (16.2) and met-enkephalin (15.8) is shown. **d**) The presence of excess (90 µM) **368** results in decreased agonist potencies for a variety of orthosteric agonists (fentanyl, circles; met-enkephalin, squares; morphine, triangles) across a series of G_i/o_ family G-protein effectors as observed by the TRUPATH assay for G-protein activation. The calculated log(EC_50_) for all agonist/G-protein combinations are right-shifted by ∼1-1.5 units in the presence of **368**. Error bars correspond to the additive fitted error 95% confidence intervals for EC_50_ values with and without **368**. **e**) This G_i/o_ family inhibition results in dampened cAMP inhibition in cells with morphine (dark blue vs. pink) and met-enkephalin (green vs. purple). Titration of **368** at EC_80_ concentrations of orthosteric agonists results in the reversal of cAMP inhibition, though with weaker potencies than those observed biochemically.

In summary, biochemical and in-cell studies suggest that **368** acts as an inhibitor of µOR-stimulated G-protein activation by several distinct orthosteric agonists. Because **368** enhances the affinity of orthosteric antagonist naloxone (with an EC_50_ ∼ 133 nM) it suggests an allosteric mechanism of µOR inhibition, though lower potencies observed in-cell assays (Fig. 3e) in the absence of naloxone hint at weaker binding of the NAM to active states of µOR.

### Structural mechanism of negative allostery at the µOR

No potent, specific NAMs exist for the µOR, though some ligands for other receptors (including CB1) can act as weak NAMs at µOR^13,25^. As **368** appears to have these desirable properties, we wanted a detailed structural characterization of its mechanism of action. To that end, we bound Nb6, an inactive κOR-specific nanobody^26,27^, to a µOR/κOR ICL3 chimera in the presence of naloxone and **368**, following established methodology for inactive GPCR structure determination^28^ (Fig. S2a). The complex was purified on FLAG resin (Fig. S2a) and subjected to cryoEM imaging (Fig. S2b). A final density map at 3.26 Å nominal resolution (Fig. S2b, Table S5) was then used to unambiguously place naloxone in the orthosteric pocket (Fig. 4a, Fig. S2b). We also observed density adjacent to naloxone, not previously observed in other inactive µOR structures (Fig. 4a, Fig. S2b)^28,29^. While the initial hit for **368** was a racemic mixture due to a single stereocenter, DEL enrichment data suggested better binding for the *S*- isomer (Fig. S3a). We designed a synthesis strategy to generate a large scale of enantiomerically pure versions of **368** (described in detail in the *Materials and Methods*, Schemes 1-3). A multi-gram scale synthesis of (*S*)-**368** as well as (±)-**368** was also developed. Using this enantiomerically pure material, we demonstrated that the *S*-isomer was the active species by both radioligand binding (Fig. S3b) and GTP depletion (Fig. S3c) assays. Thus, (*S)*-**368** was placed and docked into the final density map^30^. The resulting structure without Nb6 was also simulated for 200 ns with minimal movement of naloxone and **368**, consistent with stable poses for both molecules (Fig. S3d, S3e), though some alternate conformations within the binding pocket are observed for **368**. Finally, to support the placement of **368**, we added steric bulk to several smaller, hydrophobic residues contacting the predicted **368** binding site to introduce clash with the NAM. Both mutations (A323L and I71W) decrease the observed affinity of **368** for enhancing ^3^H-naloxone binding (Fig. S3f) and A323L substantially diminishes the extent of the observed improvement in ^3^H-naloxone affinity (Fig. S3f).

**Figure 4.**
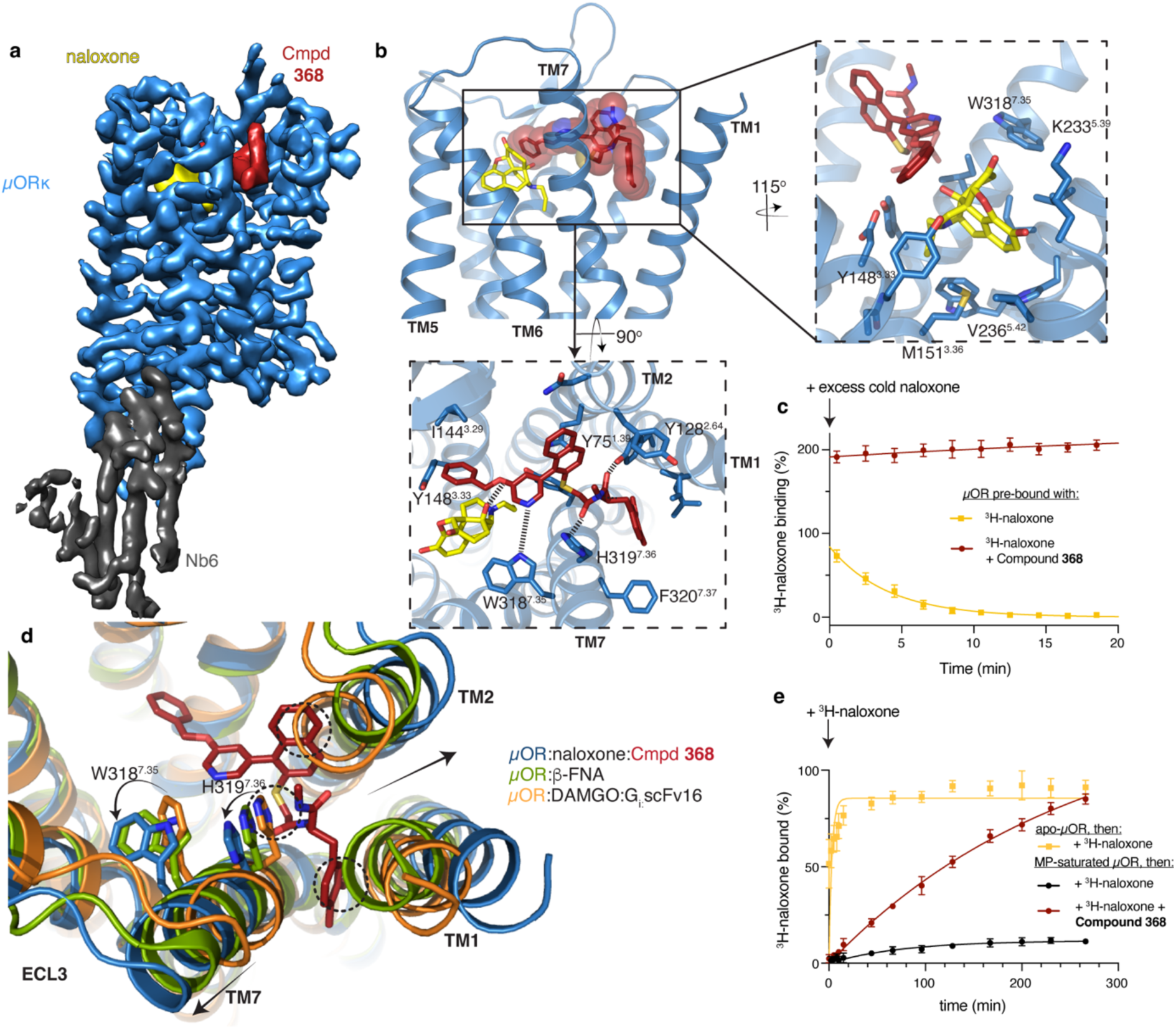
Structural mechanism of Cmpd 368 NAM activity. **a**) Cryo-EM density map of naloxone, **368**, and Nb6-bound µORκ colored by subunit. Naloxone, yellow; **368**, red; µORκ, blue; Nb6, dark gray. (**b**) Naloxone (right panel) forms a series of inactive-state interactions with µOR observed for other morphinan antagonists, though in the present structure it also forms contacts with **368** (sticks and transparent spheres). **368** forms hydrogen bonds (dashed lines) and hydrophobic interactions across the extracellular vestibule of the receptor (bottom panel), ranging from I144^3.29^ and Y148^3.33^ in TM3, to Y128^2.64^ in TM2, to F320^7.37^ in TM7. (**c**) The off-rate of ^3^H-naloxone from µOR was measured in the absence (yellow) or presence (red) of **368** by introducing excess concentrations of cold naloxone. The half-life of ^3^H-naloxone on the receptor is 2.8 minutes, but is not measurable in the presence of **368**. Data are displayed as average percent binding, normalized to non-specific binding (0%) and values in the absence of introduction of cold naloxone (100%) with error bars representing the standard deviation. (**d**) Comparing the present **368** and naloxone bound inactive state of the µOR with the active, DAMGO/G_i_-bound state of the receptor (orange, PDB:) structure and previous inactive (green) structures reveals several features about the inactivation mechanism of **368**. The pose of **368** observed here is inconsistent with either active or inactive inwards states of TM1 and TM2, resulting in a significant (though likely not functional) outward movement in both (clashes highlighted in black dashed circles). Most significantly, the active state of TM7, including W318^7.35^ and H319^7.36^, clashes extensively with **368**. Binding of **368** then likely stabilizes the inactive state of the extracellular half of TM7, resulting in an even further outward conformation compared to the previous inactive structure. (**e**) ^3^H-naloxone on-rate experiment to agonist (MP) saturated µOR, demonstrating that **368** (100 µM) can bind allosterically with the agonist and increase its observed off-rate.

Naloxone in the orthosteric binding site and makes a series of contacts very similar to those observed for the covalent antagonist β-FNA (Fig. 4b, right panel)^29^. **368** binds in the extracellular vestibule region in direct proximity to naloxone (Fig. 4b). Hydrophobic contacts are present across the terminal benzene ring in the third fragment of **368** and naloxone, as well as a stabilizing hydrogen bond between the two molecules (Fig. 4b, bottom panel). Physical contact with naloxone, as well as a slight restriction in access to extracellular solution (Fig. S4a), may explain its enhanced affinity in the presence of **368** (Fig. 2b); it also directly impacts ligand kinetics. ^3^H-naloxone has a half-life on purified µOR of 2.8 minutes (Fig. 4c). In the presence of **368**, the observed ^3^H-naloxone binding is elevated, consistent with previous results demonstrating the NAM enhancement of naloxone affinity for the receptor (Fig. 2b, S1b). Further, no observed decrease in ^3^H-naloxone binding occurred over the course of the experiment (∼30 minutes), demonstrating that this enhanced affinity is due, at least in part, to a decreased off-rate from the receptor, consistent with structural predictions (Fig. S4a). Overlay of various active, agonistb-ound structures with **368**-bound receptor suggests little to no direct clash with small molecule agonists (Fig. S4b, top) and extensive clash with peptide agonists (Fig. S4b, bottom). **368** may then act more like a competitive antagonist for endogenous opioids and a NAM for small molecule opioids. Indeed, 2 µM **368** is sufficient to fully inhibit fentanyl-induced µOR GTP turnover but has a much smaller effect on met-enkephalin-induced turnover at this concentration (Fig. S4c).

We next tested the ability of **368** to inhibit signaling by other opioid receptor subtypes (µOR, *δ*OR and κOR) using the TRUPATH assay. Increasing concentrations of **368** (up to 30 µM) results in a 3.7- fold right-shift in DAMGO potency at the µOR (Fig. S4d, Table S6), consistent with its activity previously described. The NAM also causes a right-shift in DPDPE potency at *δ*OR, though to a lesser extent (2.9- fold) (Fig. S4e, Table S6). No effect is observed at κOR activation by U50,488 (Fig. S4f, Table S6). Sequence alignment of key regions of the receptor interacting with **368** show that there is extensive conservation in most areas of the binding site (Fig. S4g) with some key exceptions, including an area at the beginning of TM7 including several aromatic residues (W318^7.35^ and H319^7.36^) predicted to make key interactions with **368** in the µOR.

**368** does not act solely by favorable contacts with bound orthosteric naloxone (Fig. 4b, bottom panel). It also makes a series of inactive-state specific interactions across the entire vestibule, ranging from the orthosteric site to residues in TM1, TM2, and TM7. A notable feature of the **368**-bound µOR is the extreme outward movement of TM1, a conformation forced by the “intercalation” of the terminal methoxybenzyl moiety in the NAM between TM1 and TM7 (Fig. 4d), though this further outward vestibule conformation appears not correlated with the receptor activation state. The naphthalene ring in **368** is in a position that would result in extensive steric clash with the extracellular half of TM2 in the DAMGO/G_i_-bound active state^31^ of the receptor (Fig. 4d). NAM binding is also incompatible with the TM2 conformation observed in the previous inactive, β-FNA-bound structure (Fig. 4d),^29^ resulting in a further outward movement of the helix compared with previous active and inactive conformations of TM2. The nitrogen in the pyridine ring of **368** is hydrogen bonded with W318^7.35^ in TM7, an interaction that can only be formed in the inactive conformation of TM7, as W318^7.35^ in the active DAMGO-bound state of the receptor is in direct clash with the NAM (Fig. 4d). Further, H319^7.36^ is hydrogen bonded with the “backbone” carbonyl and amide in **368** in the present structure, though H319^7.36^ in both the active, agonist and G-protein bound as well as the antagonist bound structure are both in steric clash with **368** (Fig. 4d). These TM7 interactions with the NAM appear to force the extracellular half of TM7 into an even further outward state as compared with previous structures, potentially consistent with the observed biochemical activity of **368** as an inverse-ago NAM (Fig. 2d) rather than a neutral antagonist. Given this series of inactive-state specific interactions at residues allosteric to the conventional opioid binding site, we speculated that **368** may modulate off-rates of opioid agonists. We saturated purified µOR with potent partial agonist mitragynine pseudoindoxyl (MP)^32^ and observed agonist off-rate by adding ^3^H-naloxone in the presence or absence of excess **368** (Fig. 4e). ^3^H-naloxone very slowly binds to the receptor, presumably due to the high potency and low off-rate of MP (Fig. 4e). However, **368** can bind MP-bound receptor and dramatically increase the off-rate of MP, suggesting that it may interact with the receptor contemporaneously with and destabilize the interactions of orthosteric agonist (Fig. 4e), though this effect may also be possible due to the modulations in naloxone off-rate explored above (Fig. 4c). Finally, selective binding of the NAM to this structurally distinct inactive-state of the receptor may explain the reduced potency of the modulator in agonist-only experiments (Fig. 2a, Fig. 3e, Fig. S4d) and suggests that the extensive cooperativity observed between naloxone and **368** may be taken advantage of *in vivo*.

### 368 boosts the ability of naloxone to reverse the effects of morphine in vivo

Given the substantial effects of the NAM on naloxone affinity and kinetics, an important measure because molecules that increase the residence time of naloxone on the receptor may have utility in reversing opioid overdoses, we next attempted to characterize its ability to potentiate naloxone *in vivo* using mouse models. When administered intravenously at 10 mg/kg in mice, **368** enters the brain with moderate penetration (∼25% of maximum plasma levels, corresponding to >10-fold the affinity of **368**) with a reasonable observed half-life (0.74 hours) (Fig. S5a). We first tested the ability of **368** to potentiate low-dose naloxone reversal of morphine induced antinociception. While naloxone is typically administered at doses of 1 or 10 mg/kg, s.c. to efficiently inhibit morphine-induced antinociception^33–35^, the dose used here (0.1 mg/kg s.c.) was confirmed to have no significant antagonism (Fig. 5a). While **368** alone has no significant impact on morphine-induced antinociception (Fig. S5b), in the presence of low-dose naloxone, **368** significantly inhibits morphine-induced antinociception in a dose-dependent manner (Fig. 5a). The median dose of **368** to potentiate low-dose naloxone antagonism of morphine antinociception was 1.15 (0.03-4.04) mg/kg, s.c. We next tested the ability of **368**, in conjunction with the low (0.1 mg/kg) dose of naloxone, to inhibit morphine-mediated respiratory depression and hyperlocomotion using the Comprehensive Lab Animal Monitoring System (CLAMS) assay.^11,36^ Morphine (20 mg/kg, s.c.) significantly increased ambulations for over 140 min (Fig. 5b) and decreased breathing rate through 100 min (Fig. 5c). Pretreatment with the low dose of naloxone was ineffective in ameliorating morphine-induced hyperlocomotion (Fig. 5b), but significantly (if modestly) reduced morphine-induced respiratory depression (Fig. 5c). While **368** itself does not change ambulation (Fig. S5c) or breathing rate relative to vehicle (Fig. S5d), it significantly enhances low-dose naloxone inhibition of hyperlocomotion (Fig. 5b) and respiratory depression (Fig. 5c) induced by morphine, in a dose-dependent manner maximal at the 100 mg/kg dose. Curiously, although **368** substantially potentiates naloxone antagonism of morphine-induced antinociception and respiratory depression, this effect was not observed against fentanyl (0.4 mg/kg)-induced antinociception (Fig. S5e). This may be due to either the probe dependence of **368** for morphinan ligands, or its low potency relative to fentanyl. Future chemical optimization and discovery efforts will focus on developing NAMs capable of inhibiting fentanyl activity *in vivo*.

**Figure 5.**
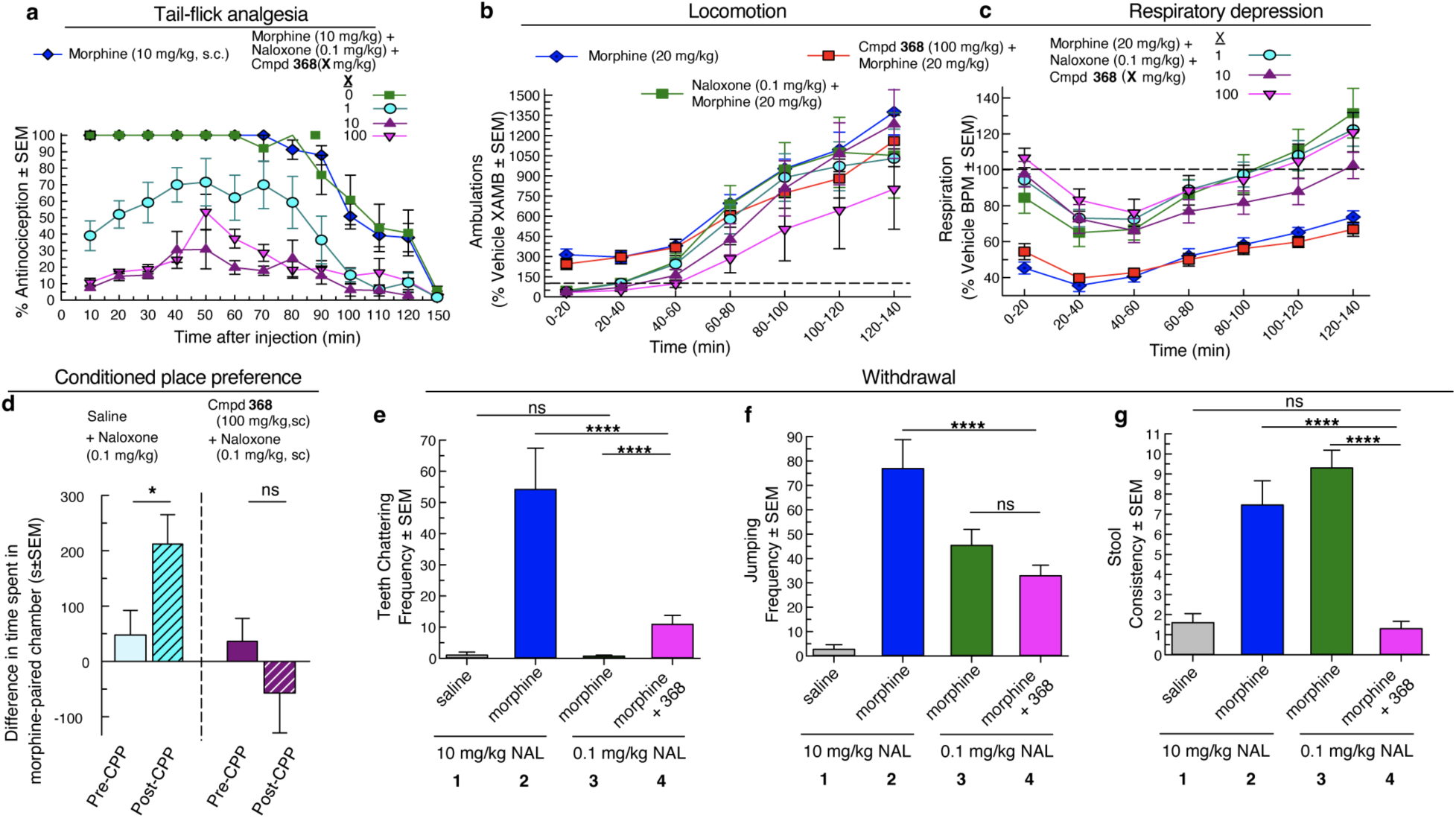
368 potentiates naloxone activity *in vivo*. (**a**) Antinociceptive time course experiment demonstrating that while low-doses of naloxone (0.1 mg/kg s.c.) have no impact on morphine (10 mg/kg) induced antinociception (green vs. blue; *F*_(12,168)_=0.49, p=0.92; Two-way RM ANOVA), increasing doses of **368** (1 mg/kg, cyan; 10 mg/kg, purple; 100 mg/kg, pink) ahead of morphine treatment in the presence of this low-dose of naloxone results in substantial inhibition of morphine-induced antinociception (*F*_(4,35)_=50.1, p<0.0001; Two-way RM ANOVA). Data are displayed as the percentage antinociception ± s.e.m. (**b,c**) Morphine (20 mg/kg s.c.) results in significantly enhanced locomotion (*F*_(6,108)_=54.7, p<0.0001; Two-way RM ANOVA with Sidak post hoc test) and decreased breathing rate (*F*_(6,108)_=4.09, p=0.001; Two-way RM ANOVA). Low doses of naloxone (0.1 mg/kg, s.c.) was unable to inhibit hyperlocomotion (*F*_(1,18)_=0.62, p=0.44; Two-way RM ANOVA) yet modestly inhibited respiratory depression (*F*_(1,18)_=11.7, p=0.003; Two-way RM ANOVA). While **368** alone has no significant impact on morphine-induced hyperlocomotion (**b**) (*F*_(6,108)_=0.36, p=0.90; Two-way RM ANOVA) or respiratory depression (**c**) (*F*_(6,108)_=1.19, p=0.32; Two-way RM ANOVA), a similar dose-dependent significant enhancement of low-dose naloxone effects on hyperlocomotion (*F*_(24,306)_=4.27, p<0.0001; Two-way RM ANOVA with Tukey post hoc test) and respiratory depression (*F*_(24,294)_=5.00, p<0.0001; Two-way RM ANOVA with Tukey post hoc test) is observed. Data are displayed in 20-minute bins as percentages relative to vehicle responses ± s.e.m. While low-dose naloxone (0.1 mg/kg) alone allows for significant (p=0.02; Student’s t-test) morphine-induced CPP (left bars) (**d**), pre-treatment with **368** (100 mg/kg) eliminates this place preference (right bars) (p=0.27; Student’s t-test). (**e,f,g**) Naloxone precipitation of withdrawal symptoms were then measured comparing conventional naloxone induced effects (**2**) with low-doses used above without (**3**) and with (**4**) pre-treatment with **368**. Various responses including teeth chattering frequency (**e**), jumping frequency (**f**) and stool consistency (**g**) were measured. (**e**) **368** with low-dose naloxone increases teeth chattering frequency relative to low-dose naloxone alone, though not to the same extent as full dose naloxone (*F*_(3,37)_= 12.8, p<0.0001; One-way ANOVA with Tukey post hoc test). (**f**) Low-dose naloxone alone causes substantial increases in jumping frequency (F_(3,37)_=17.0, p<0.0001; One-way ANOVA) that is not enhanced by **368** (P=0.66, Tukey post hoc test). (**g**) Low and high dose naloxone causes significant diarrhea in mice (F_(3,37)_=23.8, p<0.0001; One-way ANOVA with Tukey post hoc test), though this response is inhibited by **368** such that there is no difference from saline control (P=0.61, Tukey’s post hoc test). The data are displayed as average values ± s.e.m. P values are denoted in panels **d-g** as follows: ns (P>0.05), * (P≤0.05), ** (P≤0.01), *** (P≤0.001), and **** (P≤0.0001).

We next tested the combined effects of the NAM and a low-dose (0.1 mg/kg, s.c.) of naloxone for inhibition of morphine-conditioned place preference (CPP).^11,36^ While pretreatment of mice with vehicle (-30 min) and the low dose of naloxone (-15 min) alone did not prevent significant morphine CPP (Fig. 5d, left pair of bars), pretreatment with **368** (100 mg/kg, sc, -30 min) potentiated the MOR antagonism produced by the low dose of naloxone to effectively eliminate morphine CPP (Fig. 5d, right pair of bars).

Finally, as **368** treatment potentiates low-doses of naloxone in reversing morphine effects in a wide array of assays (Fig. 5a, 5b, 5c, 5d), we examined if **368** pretreatment potentiates naloxone-precipitated withdrawal symptoms^37,38^, assessing if those observed following administration of **368** (100 mg/kg, s.c.) and the low-dose of naloxone (0.1 mg/kg, s.c.) compare with the symptoms precipitated by sole treatment with conventional higher doses of naloxone (10 mg/kg, s.c.) alone. To that end, mice were first repeatedly administered saline (control group **1**), or morphine at escalating doses (10-75 mg/kg, i.p.) over the course of 5 days as utilized previously^37,38^. Assessment of withdrawal symptoms on the fifth day followed treatment with either the conventional high dose of naloxone, (10 mg/kg, group **2**), the low-dose of naloxone used for experiments herein (0.1 mg/kg, group **3**), or the low-dose of naloxone following treatment with **368** (100 mg/kg, group **4**) (Fig. 5e, 5f, 5g). The battery of withdrawal-associated behaviors following naloxone treatment (summarized in Table S7) demonstrated variable responses, but surprisingly, the combined low-dose naloxone/NAM treatment does not potentiate the magnitude of withdrawal precipitated by a low-dose of naloxone alone to the same levels of withdrawal symptoms as the conventional naloxone treatment. For some behaviors (e.g. teeth chattering frequency, Fig. 5e), low-dose naloxone treatment alone has no effect relative to saline alone (P=0.76), and the addition of **368** increases withdrawal symptoms relative to both (P<0.0001, Tukey’s post hoc test following one-way ANOVA), though notably significantly less than the response to high-dose naloxone treatment (P<0.0001). For other measures, such as jumping frequency, low-dose naloxone treatment alone precipitates significant withdrawal symptoms (Fig. 5f), but which is not significantly altered in the presence of **368** (P=0.66). Of interest, while low- and high-dose naloxone treatment precipitates significant diarrhea in morphine-dependent mice (Fig. 5g), low-dose naloxone in combination with **368** treatment does not significantly change stool consistency relative to saline control (P=0.61).

## Discussion

Allosteric modulation of GPCR activity remains a promising pathway to therapeutics that work with endogenous signaling molecules, decreasing side effects arising from both off-target effects and off-pathway activation events (e.g. biased signaling).^39–41^ Recent structures of allosteric modulator-bound Family A & B GPCRs have highlighted the diversity of potential binding sites as well as the resulting effects on receptor activation pathway(s).^21,40,42–45^ The µOR is an exception; the total lack of NAM scaffolds targeting the µOR represents a largely unexploited area of both study and potential therapeutic intervention, especially given the current opioid overdose epidemic.

We have shown that DEL hits from this screening strategy can be useful compounds for investigating mechanisms of negative allosteric modulation of the µOR as well as then lead directly to useful *in vivo* properties. We show that **368** acts as a NAM by both enhancing naloxone binding to the receptor (increasing its affinity and decreasing its off-rate) and independently stabilizing an inactive conformation of the extracellular vestibule including TM7 and ECL3. This allosteric inhibition by **368** increases the off-rate of orthosteric agonists. While **368** has minimal effect on its own in mouse behavioral assays, it potentiates naloxone-induced inhibition of a variety of morphine effects, including antinociception, hyperlocomotion and respiratory depression (Fig. 5). It can enhance the antagonism of low-dose naloxone while not resulting in the same extent of withdrawal effects observed for conventional, high-dose naloxone treatment (Fig. 5e, 5f, 5g). Future development of **368** will focus on expanding this “naloxone-boosting” effect to include inhibition of even highly potent opioid (e.g. fentanyl) effects.

The targeted screening strategy used herein can potentially be used to generate negative small molecule allosteric modulators of other GPCRs. Our selection strategy is not just based on the *presence* of allosteric pockets^46^ but necessitates their *change* in structure upon a change in the activation state of the receptor. In other words, the strategy targets sites that are strongly correlated with receptor activation and signaling. The potential challenges of the molecules discovered here present opportunities for further selections. Future studies should anticipate the probe dependence we observed for **368** and may even be able to select for modulators that are specific for orthosteric molecules of interest. Finally, our selection methodology allows the detection of molecules that bind to any conformation that is present within the active or inactive ensembles, as evidenced by the distinct inactive conformation of the receptor as compared to previous antagonist-bound structures^28,29^ (Fig. 4d). Selecting for ensembles of states with DEL screens is a way of exploiting this intrinsic plasticity of proteins, especially GPCRs.

## Acknowledgements

We thank WuXi AppTec for providing the DEL and off-DNA synthesis of hits as well as J. Su from the DELopen team for providing SAR information for chemical families. We thank M. Robertson for providing the µORκ and Nb6 vectors. Cryo-EM data were collected at S2C2. K.K. was supported by the American Diabetes Association (ADA) Postdoctoral Fellowship. E.S.O. was supported by the American Heart Association (AHA) Postdoctoral Fellowship. B.K.K. was supported by the Chan Zuckerberg Biohub.

## Author Contributions

E.S.O., J.P.M. & B.K.K. wrote the manuscript with input from all authors. E.S.O, W.H., K.K. & B.K.K. designed the DEL screening strategy. E.S.O. performed DEL selections. B.W. & E.S.O. designed, optimized, and performed radioligand binding experiments. E.S.O., W.H. & Y.S. optimized and performed GTP turnover assays. J.M.M. performed in-cell cAMP experiments. E.S.O. formed complexes for cryo-EM studies, collected and processed cryo-EM data with assistance from H.W. and C.Z., built the structural models with assistance from K.K., and performed and analyzed MD simulations. V.R. synthesized Cmpd **368** pure enantiomers and developed a scale up synthesis of racemic Cmpd **368** under S.M.’s supervision. K.A. carried out PK analysis of Cmpd **368** under S.M.’s supervision. A.E. performed TRUPATH experiments with supervision from T.C. S.O.E & J.P.M. performed and analyzed all behavioral pharmacology experiments.

## Competing Interests

B.K.K. is a founder and consultant for ConfometRx. S.M. is a founder of Sparian Biosciences. E.S.O, K.K., and B.K.K. have filed a patent around the new modulator compounds acting through µOR. The authors declare no other competing interests.

**Correspondence and requests for materials** should be addressed to Jay P. McLaughlin, Susruta Majumdar or Brian K. Kobilka.

## Supplementary Information

### Purification of µ-opioid receptor and variants

#### Purification of mouse µ-opioid receptor from sf9 cells

The mouse µ-opioid receptor was grown and purified as previously described^31^. Mouse µ-opioid receptor (µOR) with an N-terminal FLAG and C-terminal hexa-histidine tag was expressed as previously described^31^ using the baculovirus method in Spodoptera frugiperda (Sf9) cells. Naloxone was added to 10 µM final concentration upon infection and cells were collected 48 hours post-infection and stored at -80°C for later purification. µOR was extracted from membranes with 0.8% *n*-dodecyl-ß-D-malopyranoside (DDM; Anatrace), 0.08% cholesterol hemisuccinate (CHS), and 0.3% 3-((3-cholamidopropyl) dimethylammonio)-1-propanesulfonate (CHAPS; Anatrace) in 20 mM HEPES pH 7.5, 500 mM sodium chloride (NaCl), 30% glycerol, 5 mM imidazole, 10 µM naloxone, and the protease inhibitors benzamidine and leupeptin, along with benzonase (Sigma-Aldrich) to degrade cellular DNA. Cells were dounced 30 times on ice and the membranes were allowed to solubilize in the detergent for 2 hours with stirring at 4 °C, followed by centrifugation for 40 minutes at 14k rpm to pellet cell debris. The supernatant was applied to nickel-chelating sepharose resin and bound to resin with end-over-end shaking for 2 hours at 4°C. The resin was then batch washed 4 times followed by washing with 10 column volumes on-column with nickel wash buffer composed of 20 mM HEPES pH 7.4, 500 mM NaCl, 0.1% DDM, 0.01% CHS, 0.03% CHAPS, 5 mM imidazole, 10 µM naloxone, and protease inhibitors leupeptin and benzamidine. The nickel-pure µOR was then eluted in the same buffer with 250 mM imidazole. The nickel elution was initially exchanged to lauryl maltose neopentyl glycol (L-MNG; Anatrace) detergent by incubating with 0.5% L-MNG, 0.17% glycol-diosgenin (GDN; Anatrace) and 0.067% CHS overnight at 4°C. 2 mM calcium chloride (CaCl_2_) was then added and subsequently applied to M1 anti-FLAG immunoaffinity resin. The M1-bound receptor was first washed with 20 mM HEPES pH 7.4, 500 mM NaCl, 0.1% MNG, 0.033% GDN, 0.0133% CHS, 2 mM CaCl_2_, and 10 µM naloxone. The protein was then washed with 10 column volumes of 20 mM HEPES pH 7.4, 100 mM NaCl, 0.005% MNG, 0.0017% GDN, 0.00067% CHS and 2 mM CaCl_2_ followed by elution with 20 mM HEPES pH 7.4, 100 mM NaCl, 0.003% MNG, 0.001% GDN, 0.0004% CHS, 5 mM ethylenediaminetetraacetic acid (EDTA) and FLAG peptide. Multimers and dimers of the receptor were removed with size exclusion chromatography on an S200 10/300 Increase gel filtration column (GE Healthcare) equilibrated with 20 mM HEPES pH 7.4, 100 mM NaCl, 0.003% MNG, 0.001% GDN, 0.0004% CHS. Pure, monomeric apo µOR was spin concentrated to ∼150 µM, flash frozen in liquid nitrogen and stored at -80°C until further use.

The µOR-κ_ICL3_ used for the inactive state structural determination was purified in an identical fashion to the mouse µOR above. Two points mutations in ICL3 (M264L and K269R) are present in the construct to allow for binding to the κ opioid receptor (κOR)-specific Nb6 as described by Robertson et. al.^28^.

#### Purification of human µ-opioid receptor from Expi293 cells

The gene for the full-length human µ-opioid receptor was cloned into a vector for inducible expression in Expi293F cells (Thermo Fisher) with N-terminal HA signal peptide and FLAG tags and a C-terminal hexahistidine tag. This construct was transfected into Expi293F cells constitutively expressing the tetracycline repressor (Thermo Fisher) with the Expifectamine transfection kit (Thermo Fisher) following manufacturers directions with induction of receptor expression 2 days after transfection with doxycycline (4 µg/mL and 5 mM sodium butyrate) in the presence of 10 µM naloxone. Pellets were collected and frozen at -80 °C ∼30 hours after induction for later protein purification. Cells were dounced and membranes solubilized with 20 mM HEPES pH 7.5, 100 mM NaCl, 20% glycerol, 1% L-MNG, 0.1% CHS, 10 µM naloxone, protease inhibitors leupeptin and benzamidine, and benzonase, followed by purification on anti-FLAG immunoaffinity resin as above. Multimers and dimers of the receptor were removed with size exclusion chromatography on an S200 10/300 Increase gel filtration column (GE Healthcare) equilibrated with 20 mM HEPES pH 7.4, 100 mM NaCl, 0.003% MNG, 0.001% GDN, 0.0004% CHS. Pure, monomeric apo µOR was spin concentrated to ∼150 µM, flash frozen in liquid nitrogen and stored at -80°C until further use.

### Expression & purification of heterotrimeric G-proteins

Heterotrimeric G_i_ was expressed in *Trichoplusia ni* (*T. ni*) with the BestBac method (Expression Systems) and purified as previously described.^31,47^ Briefly, T. ni cells were infected with one virus encoding the wild-type human Gα_i_ subunit and another encoding the wild-type human β_1_γ_2_ subunits with a histidine tag inserted at the N-terminus of the β subunit. Cells were harvested 48 hours post infection and lysed with hypotonic buffer. Heterotrimeric Gα_i_β_1_γ_2_ proteins were extracted in a buffer containing 1% sodium cholate and 0.05% DDM. The heterotrimer was purified using nickel-chelating sepharose chromatography while removing cholate. Human rhinovirus 3C protease (3C protease) was added to cleave off the histidine tag overnight at 4°C on-column. The flow through was collected and dephosphorylated with lambda protein phosphatase (NEB), calf intestinal phosphatase (NEB) and Antarctic phosphatase (NEB) in the presence of 1 mM manganese chloride (MnCl_2_). The heterotrimer was further purified by ion exchange chromatography on a MonoQ 10/100 GL column (GE Healthcare) in 20 mM HEPES pH 7.5, 1 mM MgCl_2_, 0.05% DDM, 100 µM TCEP, and 20 µM GDP and eluted with a linear NaCl gradient from 50 to 500 mM. The purified heterotrimer was collected and dialyzed into 20 mM HEPES pH 7.5, 100 mM NaCl, 0.02% DDM, 100 µM TCEP and 20 µM GDP overnight at 4°C, followed by concentration to <250 µM, addition of 20% glycerol and flash-freezing in liquid nitrogen and storage at -80°C until further use.

### Purification of Nb6

Nb6 was grown and purified in a similar manner to that previously described^28^. Briefly, BL21 *Escherichia coli* were transformed with Nb6 plasmid^27^ and single colonies were picked into small scale starter cultures in LB with 100 µg/mL ampicillin, grown overnight at 37°C. The starter culture was used to inoculate larger cultures of TB supplemented with 100 µg/mL ampicillin, which were grown to an optical density_600_ = 0.7. IPTG was added to 1 mM to induce expression overnight at room temperature. Pellets were harvested and flash frozen for later purification.

Pellets were resuspended with 50 mM Tris pH 8.0, 0.5 mM EDTA and 20% w/v sucrose (TES) buffer at a ratio of 2 mL buffer per gram of pellet mass in the presence of protease inhibitor cocktail and shaken for 1 hour at 4°C. An additional 4 mL of TES buffer diluted 1:4 in water per gram of pellet was added; the mixture was shaken for an additional hour at 4°C. Cell debris was pelleted by centrifugation at 18,000 rpm for 15 minutes and the resulting supernatant was removed and bound to nickel-chelating sepharose resin in batch for 1.5 hours at 4°C with gentle shaking with 5 mM imidazole supplemented to the solution. The Nb6-bound resin was then loaded to a gravity column and washed with 20 mM HEPES pH 7.4 and 100 mM NaCl followed by washing with the same buffer with 500 mM NaCl. The Nb6 was eluted in the same high-salt buffer with 250 mM imidazole. Nickel-pure Nb6 was then dialyzed overnight into 20 mM HEPES with 100 mM NaCl, followed by final purification on size exclusion chromatography on a S200 10/300 Increase column in the same buffer. Nb6 was concentrated and flash-frozen in liquid nitrogen for further experiments.

### Preparation of samples for DEL selections

#### MOR/naloxone formation

Purified µOR (above) was incubated with excess naloxone for 1 hour at room temperature before selection experiments in 20 mM HEPES pH 7.4, 100 mM NaCl, 0.003% MNG, 0.001% GDN, 0.0004% CHS.

#### µOR/G_i_/met-enkephalin complex formation

µOR/G_i_ complex was formed in a similar manner to that described previously^31,48^. Briefly, excess met-enkephalin peptide [MedChemExpress] was added to purified µOR (above) and incubated at room temperature for 1 hour. Concurrently, 1% L-MNG/0.1% CHS was added to G_i_ purified in DDM to exchange detergent on ice for 1 hour. The two reactions were mixed for a final molar ratio of 1:1.5 µOR:G_i_ and incubated at room temperature for 1 hour. Apyrase was added and the reaction was further incubated on ice for 2 hours. 2 mM CaCl_2_ was added to the reaction before adding to M1 anti-FLAG resin for 20 minutes. The column was washed with 20 mM HEPES pH 7.4, 100 mM NaCl, 0.003% MNG, 0.001% GDN, 0.0004% CHS, 5 µM met-enkephalin, and 2 mM CaCl_2_, followed by elution in the same buffer with 5 mM EDTA replacing the CaCl_2_ and FLAG peptides. After removing excess unbound G_i_ heterotrimer, excess un-coupled µOR was removed by size exclusion chromatography with a S200 10/300 Increase column into 20 mM HEPES pH 7.4 and 100 mM NaCl, 0.003% MNG, 0.001% GDN, 0.0004% CHS, 100 µM TCEP and 1 µM met-enkephalin. µOR/G_i_/met-enkephalin complex was spin concentrated to ∼25 µM and flash-frozen in liquid nitrogen after adding 15% glycerol.

### DEL selection

#### Target preparation

The two samples produced above (µOR/naloxone and µOR/G_i_/met-enkephalin) were diluted to 2 µM final concentration in **W1**: 20 mM HEPES pH 7.4, 150 mM NaCl, 100 µM TCEP, 0.1% hsDNA (Thermo Fisher Scientific), 0.02% MNG, 0.002% CHS and 20 µM ligand (naloxone and met-enkephalin, respectively). A third blank sample (no protein control) was made with the same buffer in the absence of ligand. 300 µL of HisPur^TM^ Ni-NTA magnetic beads (Thermo Fisher Scientific) as slurry were washed with water followed by splitting into 4 batches and washing with the 4 respective buffers made above. Washing with buffer was repeated 2 additional times. 300 µL of 2 µM targets (above) and 300 µL buffer without agonist was incubated with the respective magnetic Ni-NTA resin for 30 minutes with mixing to bind targets to resin. Each of the 3 samples was then split into 3 100 µL aliquots corresponding to each of the 3 rounds of selection.

#### First round of selection

Any unbound target was removed from the 3 100 µL targets by washing with 200 µL respective wash buffer above. The 3 DEL samples (G1, G2, G3) provided by WuXi (10 µL) were resuspended with 90 µL of the respective wash buffers above for the 3 conditions and then applied to the 3 targets bound to magnetic resin and incubated with shaking at room temperature for 1.5 hours to bind to the target(s). Reactions were spun briefly to settle the resin, followed by washing all with 200 µL respective buffer above. The 3 selections were then washed again with 200 µL of the same buffers with lower detergent concentrations (0.001% MNG, 0.0001% CHS) and lower ligand concentrations (500 nM) (**W2**). The 3 selections were then washed with 200 µL of the same respective buffers in the absence of detergent and even further lower ligand concentration (100 nM) (**W3**) to minimize detergent and ligand carry-over between rounds. The resin with target (and bound library molecules) was then resuspended in 100 µL of the respective last wash buffer and heated at 95 °C for 10 minutes to elute bound library components but leave receptor bound to resin. The heat denatured mixture was spun down and the supernatant removed from the resin. 10 µL of each of the 3 selections was saved for later analysis, and 90 µL was reserved for the next round of selection.

#### Second round of selection

Another 100 µL aliquot of target-bound magnetic resin was washed with 200 µL respective **W1** buffer to remove unbound target. The 3 samples of resin were then resuspended in 90 µL respective **W1**, to which the 90 µL of reserved Round 1 DEL selection was added and incubated with shaking at room temperature for 1.5 hours. Reactions were spun briefly to settle the resin, followed by washing with 200 µL respective **W1** buffer. The 3 selections were then washed again with 200 µL of respective **W2** buffer, followed by 200 µL of respective **W3** buffer. The resin with target (and bound library molecules) was then resuspended in 100 µL respective **W3** and heated at 95 °C for 10 minutes to elute bound library components but leave receptor bound to resin. The heat denatured mixture was spun down and the supernatant removed from the resin. 50 µL of each of the 3 selections was saved for later analysis, and 50 µL was reserved for the next round of selection.

#### Third round of selection

Another 100 µL aliquot of target-bound magnetic resin was washed with 200 µL respective **W1** buffer to remove unbound target. The 3 samples of resin were then resuspended in 50 µL respective **W1**, to which the 50 µL of reserved Round 2 DEL selection was added and incubated with shaking at room temperature for 1.5 hours. Reactions were spun briefly to settle the resin, followed by washing with 200 µL respective **W1** buffer. The 3 selections were then washed again with 200 µL of respective **W2** buffer, followed by 200 µL of respective **W3** buffer. The resin with target (and bound library molecules) was then resuspended in 100 µL respective **W3** and heated at 95 °C for 10 minutes to elute bound library components but leave receptor bound to resin. The heat denatured mixture was spun down and the supernatant removed from the resin. All of the final (third) round of selection supernatant for all 3 conditions was saved for subsequent sequencing and enrichment analysis.

### Preparation of µ-opioid receptor membranes

#### Insect cell membranes

Cell membranes containing the mouse µOR described above were generated by infecting Sf9 cells in an identical manner to that described above for protein purification. Sf9 cells expressing µOR were resuspended in cold lysis buffer composed of 10 mM HEPES pH 7.4, 10 mM MgCl_2_, and 20 mM potassium chloride (KCl) with the protease inhibitors leupeptin, benzamidine, and cOmplete^TM^ EDTA-free protease inhibitor cocktail tablets (Sigma-Aldrich). The lysed cells were spun at 45k rpm for 45 minutes to pellet membranes. The supernatant was removed and membrane pellets were resuspended in cold lysis buffer followed by douncing ∼30 times on ice. The dounced membranes were spun again at 45k rpm for 45 minutes. The pellets were resuspended in cold lysis buffer again with the addition of benzonase and dounced a further ∼30 times, followed by a further spin at 45k rpm for 45 minutes. Membranes were resuspended in the same lysis buffer above in the presence of 1 M NaCl and dounced a further ∼30 times and pelleted. Finally, membranes were washed with the original lysis buffer, dounced, and pelleted. Final washed membranes were resuspended to 2g original pellet mass per 1 mL in lysis buffer; the resulting µOR-containing membranes were flash frozen for later radioligand binding experiments.

### Radioligand binding experiments

#### Membrane binding experiments

µOR-containing insect cell membranes prepared above were diluted 1:1000 in 20 mM HEPES pH 7.4, 100 mM NaCl, and 0.05% bovine serum albumin (BSA). For saturation binding experiments, membranes were incubated with a serial dilution of ^3^H-naloxone (50.3 Ci/mmol; Perkin Elmer) and allosteric modulators at different constant concentrations for 1.5 hours at room temperature with shaking. For “competition” binding experiments, membranes were incubated with 2 nM ^3^H-naloxone and serially diluted allosteric modulators for 1.5 hours at room temperature with shaking. Following incubation, membranes were rapidly bound to double thick 90 x 120 mm glass fibre Printed Filtermat B filters (Perkin Elmer) and washed with cold binding buffer (20 mM HEPES pH 7.4, 100 mM NaCl) using a MicroBeta Filtermat-96 cell harvester (Perkin Elmer). ^3^H-naloxone bound membranes on Filtermats were measured with a MicroBeta counter (Perkin Elmer) after addition of MultiLex B/HS melt-on scintillator sheets (Perkin Elmer) and data values were plotted after normalizing total counts per minute to the highest and lowest values.

#### Kinetic SPA binding

Copper His-Tag PVT slurried resin (Perkin Elmer) was added 1:1 v:v with 40 mM HEPES PH 7.4, 200 mM NaCl, 0.02% MNG, 0.002% CHS, and 0.1% BSA to match buffer with the receptor. For the ^3^H-naloxone on-rate experiment (mitragynine pseudoindoxyl, MP off-rate experiment), purified µOR in detergent was diluted to 40 nM in 20 mM HEPES pH 7.4, 100 mM NaCl, 0.01% MNG, 0.001% CHS, and 0.05% BSA in the presence or absence of slight (60 nM) excess high affinity orthosteric agonist (MP). 25 µL of receptor solution was incubated with 50 µL resin slurry in buffer for 1 hour with shaking at room temperature. ^3^H-naloxone was diluted to 200 nM in 20 mM HEPES pH 7.4, 100 mM NaCl, 0.01% MNG, 0.001% CHS, and 0.05% BSA in the presence or absence of 100 µM 10105-368-30-605 NAM compound and in the presence or absence of 200 µM cold naloxone. 25 µL of the ^3^H-naloxone solutions was added to the 75 µL receptor/SPA resin mixtures to initiate radioligand binding to the receptor and the solutions were immediately read on a MicroBeta counter at various time points for ∼5 hours. Data were normalized to total counts observed for the non-specific binding condition and the maximum counts observed in the absence of MP and cold naloxone.

For the ^3^H-naloxone off-rate experiment, purified µOR in detergent was diluted to 40 nM in 20 mM HEPES pH 7.4, 100 mM NaCl, 0.01% MNG, 0.001% CHS, and 0.05% BSA with 50 nM ^3^H-naloxone in the presence or absence of 100 µM **368**. Excess cold naloxone (200 µM) was added at t=0 and counts were measured at various time points for ∼30 minutes. Counts were normalized to 0% (excess cold naloxone, no **368**) and 100% (no cold naloxone, no **368**) throughout the time course.

### Chemical Synthesis

Reagents purchased from Sigma-Aldrich Chemicals, Ambeed, Chemscene, Chemimpex and used as purchased. All reactions are performed under argon atmosphere unless otherwise specified. While performing synthesis, reaction mixtures were purified by silica gel flash chromatography on E. Merck 230–400 mesh silica gel 60 using a Teledyne ISCO CombiFlash Rf instrument with UV detection at 280 and 254 nm. RediSep Rf silica gel normal phase columns were used with a gradient range of 0–80% EtOAc in Hexane. Reported yields are isolated yields upon purification of each intermediate. Final clean (purity ≥95%, LC-MS Agilent 1100 Series LC/MSD) compounds were used for the study. NMR spectra were collected using Varian 500 MHz NMR instrument at the NMR facility of Washington University School of Medicine in St. Louis collected via the Bruker Topspin Software (Bruker Topspin 3.5 pI 6). Chemical shifts are reported in parts per million (ppm) relative to residual solvent peaks at the nearest 0.01 for proton and 0.1 for carbon (CDCl3 1H: 7.26, 13C: 77.1). Peak multiplicity is reported as the NMR spectra were processed with MestreNova software14.2.0, namely s – singlet, d – doublet, t – triplet, q – quartet, m – multiplet for examples. Coupling constant (J) values are expressed in Hz. Mass spectra were obtained at the St. Louis College of Pharmacy using the Agilent 1100 Series LC/MSD by electrospray (ESI) ionization with a gradient elution program (Ascentis Express Peptide C18 column, acetonitrile/water 5/95/95/5, 5 minutes, 0.05% formic acid) and UV detection (214 nM/254 nM). High resolution mass spectra were obtained using a Bruker 10 T APEX -Qe FTICR-MS and the accurate masses are reported for the molecular ion [M+H]+. Detailed experiments and characterization of the new compounds are included below.

#### Main synthetic scheme

**Scheme 1:**
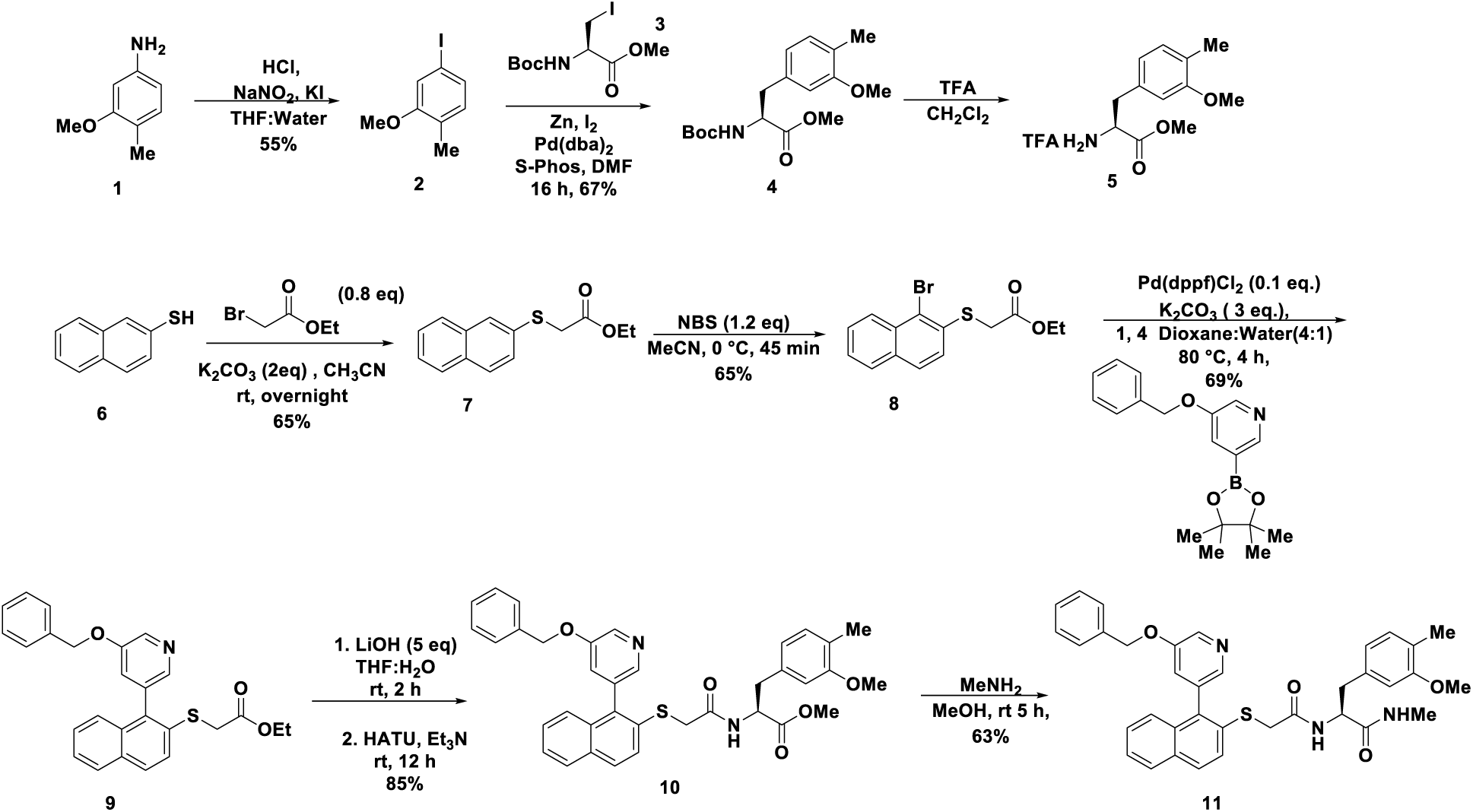
Synthesis of Cmpd (S)-368.

Synthesis of Cmpd (S)-**368** enantiomer commenced with diazotization of 3-methoxy-4-methylaniline **1** and treatment with KI gave iodo compound **2** in 85% yield. Reaction of this iodo intermediate with methyl (*R*)-2-((*tert*-butoxycarbonyl)amino)-3-iodopropanoate using Negishi coupling^49^ reaction yielded the coupling product **4** in 67%. Naphthalene-2-thiol, **6** was alkylated with ethyl bromoacetate to give compound **7** which on bromination using NBS furnished the bromo product **8** in 65% yield. Suzuki coupling^50^ on **8** on with 3-(benzyloxy)-5-(4,4,5,5-tetramethyl-1,3,2-dioxaborolan-2-yl)pyridine yielded the coupled product **9** in 69% yield. Hydrolysis of ester group in **9** gave the corresponding carboxylic acid, which was coupled with compound **5** using HATU under standard coupling conditions to give product **10** in 85% yield. An amination of ester using MeNH_2_ furnished the Cmpd **368** (*S*) **11** (Scheme 1).

**Scheme 2:**
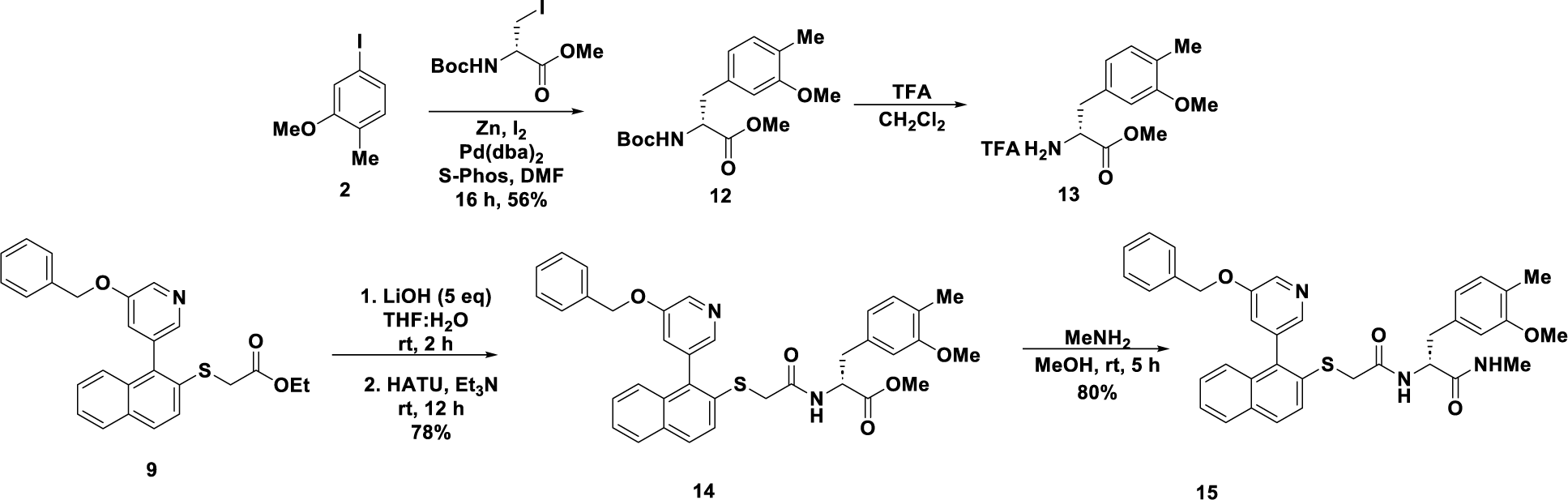
Synthesis of Cmpd 368 (*R*)

Cmpd **368** (*R*) **15** was also synthesized (Scheme 2) following a similar protocol to the (*S*) isomer.

**Scheme 3:**
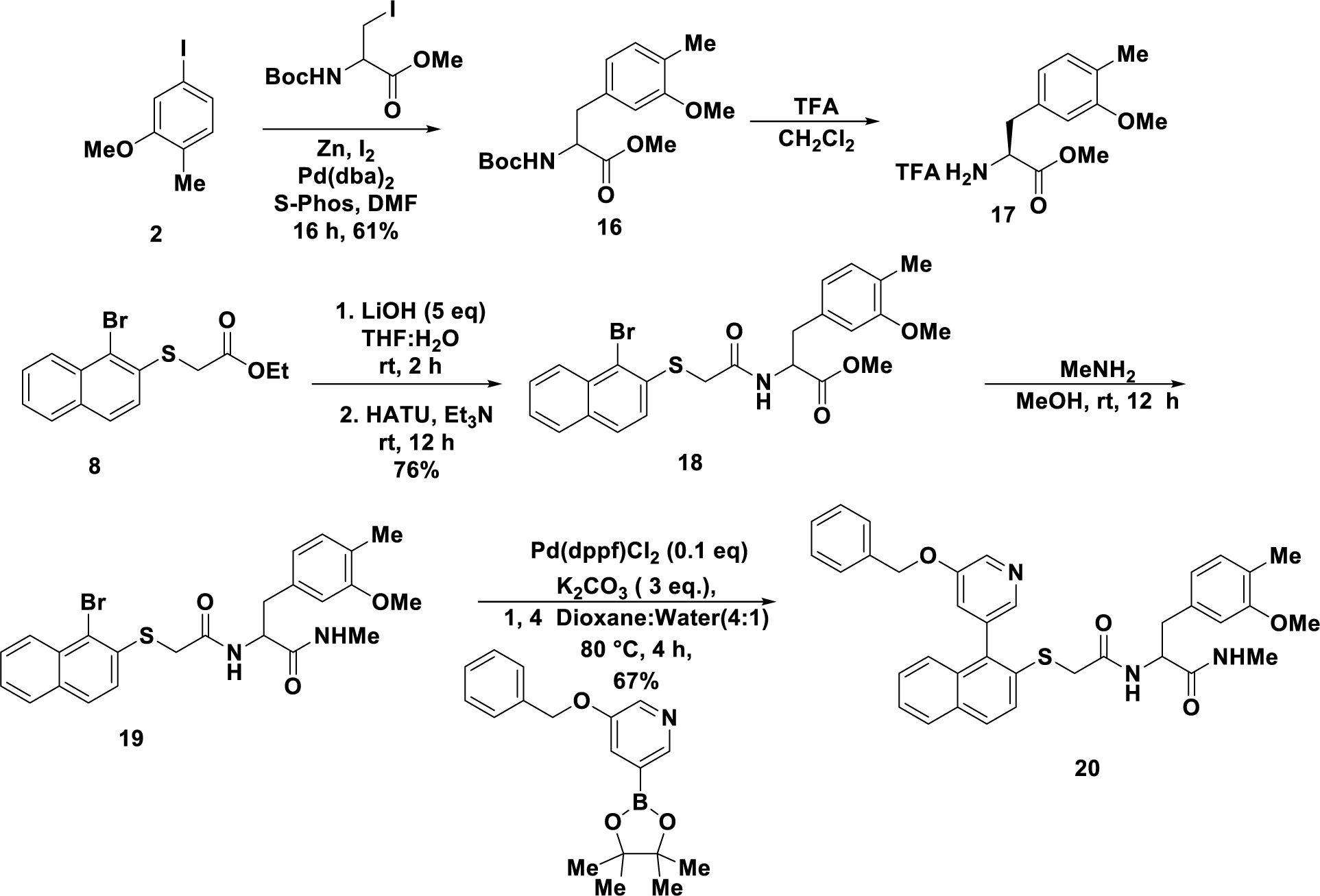
Synthesis of (±)-368.

Synthesis of (±) **368** commenced with the preparation of racemic compound **16.** Boc deprotection using trifluoracetic acid gave **17** which on coupling reaction with acid obtained from **8** furnished compound **18**. Compound **18** was subjected to an amination reaction and product obtained **19** was carried forward to the next step without purification and subjected to Suzuki reaction yielded coupling product **20** Cmpd **368** in 67% yield (Scheme 3).

#### Synthesis of precursors

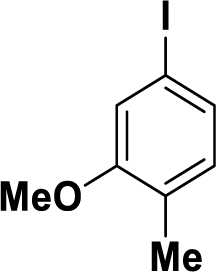

##### 4-iodo-2-methoxy-1-methylbenzene (2)

To a stirred solution of 4-iodo-2-methoxy-1-methylbenzene **1** (5 g, 36.49 mmol) in (50 mL) THF was added 15 mL HCl in (25 mL)H_2_O at 0 ^0^C. NaNO_2_ (3.02 g, 43.79 mmol) solution in (10 mL) was added dropwise and reaction mixture was stirred at the same temperature for 20 minutes. Solution of KI in (50 mL) of water was added slowly to the reaction mixture at 0 ^0^C and reaction was stirred at room temperature for another 18 hours. Reaction mixture was diluted with water and EtOAC (100 mL), the organic layer was separated and aqueous layer was extracted with EtOAc (100 mL×2). The combined organic layer was washed with saturated 10% NaOH (50 mL), and saturated Na_2_S_2_O_3_ and water. The organic layer was dried over anhydrous Na_2_SO_4_ and evaporated off and the crude reaction mixture was purified by flash chromatography using 1-5% EtOAc:Hexane yielding the iodo compound **2** (4.5 g, 50% yield) as a brown oil. ^1^H NMR (500 MHz, CDCl_3_) δ 7.16 (d, *J* = 7.7 Hz, 1H), 7.07 (s, 1H), 6.82 (d, *J* = 7.7 Hz, 1H), 3.78 (s, 3H), 2.13 (s, 3H). ^13^C NMR (126 MHz, CDCl_3_) δ 158.3, 132.0, 129.4, 126.5, 119.2, 90.4, 55.5, 15.9.

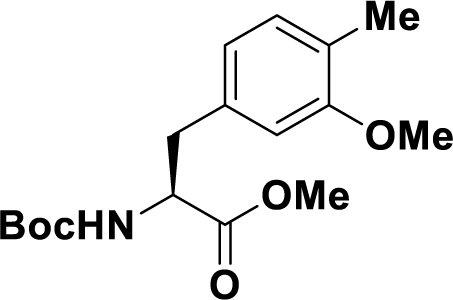

##### Methyl (*S*)-2-((*tert*-butoxycarbonyl)amino)-3-(3-methoxy-4-methylphenyl)propanoate (4)

The synthesis was carried out according to literature reports.^49^ Zinc (0.193 g, 0.80 mmol) was dissolved in DMF (2 mL) to it I_2_ (0.023 g, 0.090 mmol) was added under N_2_ (a change in color was observed from colorless to yellow and again to a colorless solution). Methyl (*R*)-2-((*tert*-butoxycarbonyl)amino)-3-iodopropanoate **3** (0.2 g, 0.60 mmol) and iodine (0.023 g, 0.090 mmol) was added. The reaction was exothermic, and the reaction was stirred at room temperature for 5 minutes. To this Pd(dba)_3_ (13.8 mg, 0.015 mmol), *S*-Phos (12 mg, 0.03 mmol) and 4-iodo-2-methoxy-1-methylbenzene (0.193 g, 0.80 mmol) were added sequentially, and reaction mixture was stirred at room temperature overnight. After completion of reaction, it was diluted with DCM (10 mL) and cold water (10 mL), the organic layer was separated, and the aqueous layer was extracted with DCM (10 mL× 2). The organic layers were combined washed with water, brine and dried over anhydrous Na_2_SO_4._ The volatile solvents were evaporated off to furnish crude residue which was purified by flash column chromatography using (5-10%, EtOAc:hexane) eventually leading to product isolation **4** (0.130 g, 67% yield). ^1^H NMR (500 MHz, CDCl_3_) δ 7.04 (d, *J* = 7.3 Hz, 1H), 6.69 – 6.55 (m, 2H), 4.97 (s, 1H), 4.63–4.51 (m, 1H), 3.81 (s, 3H), 3.73 (s, 3H), 3.17–2.94 (m, 2H), 2.18 (s, 3H), 1.43 (s, 9H). ^13^C NMR (126 MHz, CDCl_3_) δ 172.4, 156.2, 134.6, 130.6, 121.1, 110.9, 81.9, 55.2, 54.5, 52.4, 38.2, 28.3, 15.9.

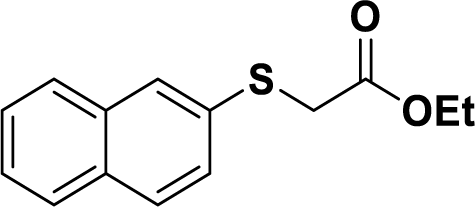

##### Ethyl 2-(naphthalen-2-ylthio)acetate (7)

Naphthalene-2-thiol **6** (1 g, 6.20 mmol) was dissolved in CH_3_CN (20 mL) at room temperature to it K_2_CO_3_ (1.72 g, 12.4 mmol) and ethyl bromoacetate (55 mL, 5 mmol) was added and reaction was stirred at room temperature overnight. To this 0.5 N NaOH (20 mL) was added and diluted with EtOAc (20 mL). The organic layer was separated, and aqueous layer was extracted with EtOAc (20 mL × 2). The organic layer was washed with brine and dried over anhydrous Na_2_SO_4._ The volatile solvents were evaporated off to furnish crude residue which was purified by flash column chromatography using (1-5%, EtOAc:hexane) yielded the product **7** (0.8 g, 65%). ^1^H NMR (500 MHz, CDCl_3_) δ 7.98 – 7.64 (m, 4H), 7.49 (ddd, *J* = 13.2, 7.0, 4.1 Hz, 3H), 4.18 (q, *J* = 7.2 Hz, 2H), 3.75 (s, 2H), 1.22 (t, *J* = 7.2 Hz, 3H).^13^C NMR (126 MHz, CDCl_3_) δ 169.6, 133.7, 132.4, 132.1, 128.6, 128.2, 127.7, 127.6, 127.3, 126.6, 126.1, 61.6, 36.7, 14.1.

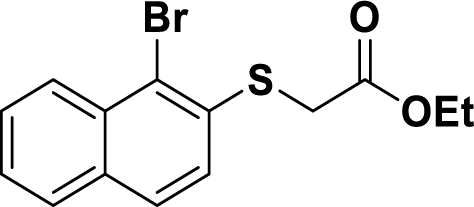

##### Ethyl 2-((1-bromonaphthalen-2-yl)thio)acetate (8)

Ethyl 2-(naphthalen-2-ylthio)acetate **7** (0.1, 0.4 mmol) was dissolved in CH_3_CN (5 mL) and cooled to 0 ^0^C, NBS (0.086 g, 0.48 mmol) was added, and the reaction stirred at same temperature for 45 minutes. After completion of reaction, it was diluted with EtOAc (20 mL), and sat. NaHCO_3,_ and water. The organic layer was separated, and aqueous layer was extracted with EtOAc (10 mL× 2). The combined organic layer was washed with brine and dried over anhydrous Na_2_SO_4_. The volatile solvents were evaporated off to furnish crude residue which was purified by flash column chromatography using (1-5%, EtOAc:hexane) yielded product **8** (0.088 g, 65% yield). ^1^H NMR (500 MHz, CDCl_3_) δ 8.24 (d, *J* = 8.5 Hz, 1H), 7.81 – 7.72 (m, 2H), 7.58 (t, *J* = 7.2 Hz, 1H), 7.53 – 7.44 (m, 2H), 4.19 (q, *J* = 7.1 Hz, 2H), 3.79 (s, 2H), 1.23 (t, *J* = 7.1 Hz, 3H). ^13^C NMR (126 MHz, CDCl_3_) δ 169.1, 134.6, 132.62,132.6 128.1, 127.9, 126.9, 126.3, 125.6, 123.3, 61.7, 35.8, 14.0.

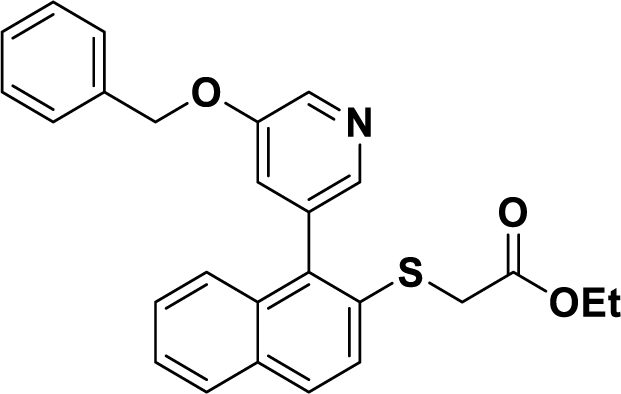

##### Ethyl 2-((1-(5-(benzyloxy)pyridin-3-yl)naphthalen-2-yl)thio)acetate (9)

Ethyl 2-((1**-**bromonaphthalen-2-yl)thio)acetate **8** (0.02 g, 0.06 mol) was dissolved in 1,4-dioxane:water (2.5 mL, 4:1) and argon was bubbled in for 10 minutes. K_2_CO_3_ (0.025 g, 0.18 mmol) and ([1,1′-Bis(diphenylphosphino)ferrocene]dichloropalladium(II)) (0.005 g, 0.001 mmol) was added and reaction mixture was stirred under argon at 80 ^0^C for 4 hours. After completion of reaction, the reaction mixture was dried over anhydrous Na_2_SO_4_ and filtered through celite pad, the celite pad washed with EtOAc (20 mL). The volatile solvents were evaporated off and the residue obtained purified by flash column chromatography to yield product **9** (0.018 g, 69% yield). ^1^H NMR (500 MHz, CDCl_3_) δ 8.52 (d, *J* = 2.3 Hz, 1H), 8.21 (s, 1H), 7.87 (dd, *J* = 12.7, 8.8 Hz, 2H), 7.65 (d, *J* = 8.7 Hz, 1H), 7.59 – 7.17 (m, 9H), 5.15 (s, 2H), 4.14 (q, *J* = 7.2 Hz, 2H), 3.55 (s, 2H) 1.20 (t, *J* = 7.1 Hz, 3H).^13^C NMR (126 MHz, CDCl_3_) δ 169.2, 154.5, 143.5, 137.7, 136.0, 135.7, 134.6, 132.8, 132.2, 132.1, 129.1, 128.6, 128.2, 128.0(2C), 127.6, 127.0(2C), 126.5, 126.0, 125.6, 123.6, 70.3, 61.5, 36.2, 14.0.

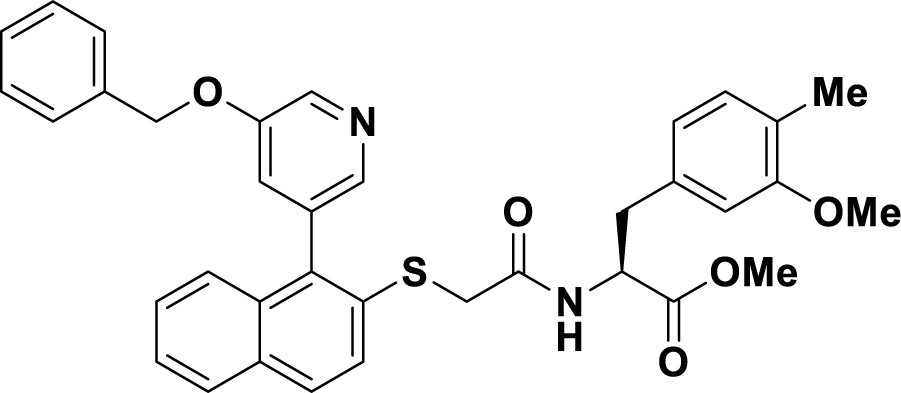

##### Methyl(*S*)-2-(2-((1-(5-(benzyloxy)pyridin-3-yl)naphthalen-2-yl)thio)acetamido)-3-(3-methoxy-4-methylphenyl)propanoate (10)

Ethyl 2-((1-(5-(benzyloxy)pyridin-3-yl)naphthalen-2-yl)thio)acetate **9** (0.05 g, 0.11 mol) was dissolved in THF:H_2_O (3 mL, 2:1), to this LiOH (1N, 0.6 mL, 0.58 mmol) was added and reaction stirred overnight. After completion of reaction, it was acidified using 1N HCl and extracted with EtOAc (20 mL× 2), the organic layer was washed with brine and dried over anhydrous Na_2_SO_4_. The volatile solvents were evaporated off to give the crude acid. Methyl (*S*)-2-((*tert*-butoxycarbonyl)amino)-3-(3-methoxy-4-methylphenyl) propanoate **4** (0.049 g, 0.15 mmol) was dissolved in CH_2_Cl_2_ (2 mL) to it TFA (0.5 mL) was added, and reaction stirred for 3 hours. After completion of reaction, the solvent was evaporated off and residue **5** dissolved in CH_2_Cl_2_:DMF (3 mL, 2:1) and to it the acid obtained above was added using DMF (1 mL) to this Et_3_N (0.05 mL, 0.35 mmol), HATU (0.067 g, 0.17 mmol) was added sequentially, and reaction mixture was stirred at room temperature overnight. The reaction mixture was diluted with cold water (10 mL) and extracted with EtOAc (20 mL× 2). The combined organic layer was washed with brine and dried over anhydrous Na_2_SO_4._ The volatile solvents were evaporated off to furnish crude residue which was purified by flash column chromatography using (1-50%, EtOAc:hexane) yielding, the product **10** (0.06 g, 85% yield). ^1^H NMR (500 MHz, CDCl_3_) δ 8.58 – 8.44 (m, 1H), 8.12 (d, *J* = 49.7 Hz, 1H), 7.91 – 7.78 (m, 2H), 7.54 – 7.25 (m, 12H), 7.04 (s, 1H), 6.39 (d, *J* = 6.9 Hz, 1H), 5.15 (s, 1H), 5.06 – 4.94 (m, 1H), 4.85 – 4.67 (m, 1H), 3.72 – 3.60 (m, 5H), 3.57 (s, 3H), 3.09 – 2.96 (m, 2H), 1.99 (m, 3H). ^13^C NMR (100 MHz, CDCl_3_) δ 171.5, 167.6, 157.7, 143.1, 135.8, 134.4, 133.9, 131.7, 130.5, 129.5, 128.7, 128.7, 128.4, 128.3, 128.2, 127.8, 127.7, 127.3, 126.0, 125.3, 125.2, 120.9, 120.6, 110.3, 70.4, 55.1, 53.4, 52.4, 37.5, 36.6, 15.7.

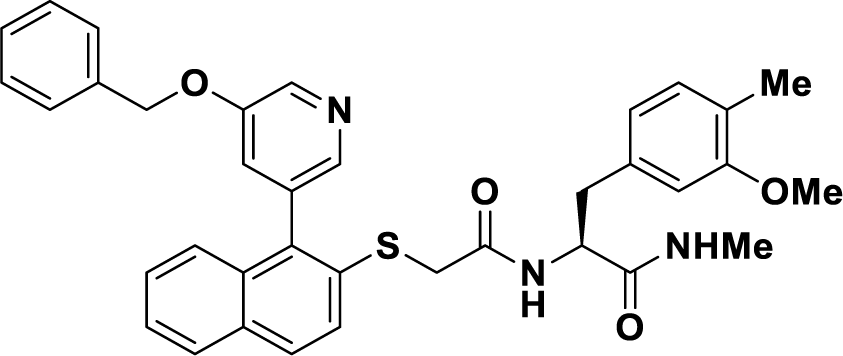

##### (*S*)-2-(2-((1-(5-(benzyloxy)pyridin-3-yl)naphthalen-2-yl)thio)acetamido)-3-(3-methoxy-4-methylphenyl)-*N*-methylpropanamide (11)

Methyl(*S*)-2-(2-((1-(5-(benzyloxy)pyridin-3-yl)naphthalen-2-yl)thio)acetamido)-3-(3-methoxy-4-methylphenyl)propanoate was dissolved in MeOH and to it MeNH_2_ (40% in H_2_O, 0.25 mL) was added and reaction stirred at room temperature overnight. After completion of reaction, volatile solvents were evaporated off and residue purified by flash chromatography using (80% EtOAc:hexane) to furnish the product **11** as a white solid (7 mg, 63% yield).^1^H NMR (500 MHz, CDCl_3_) δ 8.54 (s, 1H), 8.17 (d, *J* = 9.0 Hz, 1H), 7.92 – 7.75 (m, 2H), 7.53 – 7.26 (m, 10H), 7.12 (s, 1H), 6.79 (dd, *J* = 30.1, 7.3 Hz, 1H), 6.64 – 6.41 (m, 2H), 5.48 (d, *J* = 24.3 Hz, 1H), 5.18 (ddt, *J* = 24.7, 18.0, 9.3 Hz, 2H), 4.58 – 4.40 (m, 1H), 3.70 (s, 3H), 3.62 – 3.48 (m, 2H), 3.06 – 2.84 (m, 2H), 2.54 (s, 3H), 2.06 (s, 3H). ^13^C NMR (126 MHz, CDCl_3_) δ 170.70 (d, *J* = 4.0 Hz), 167.8, 157.9, 135.9, 135.0, 134.5, 131.9, 130.6, 129.6, 128.7, 128.3, 128.1, 127.71,127.7,127.4, 126.1, 125.4, 124.1, 124.0, 120.7, 110.6, 70.5, 55.2, 38.0, 37.2, 37.0, 36.6, 15.8. HRMS calcd for C36H35N3O4S Na+; 628.2240: HRMS found 628.2245.

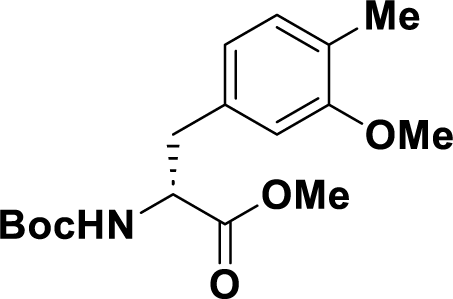

##### Methyl (*R*)-2-((*tert*-butoxycarbonyl)amino)-3-(3-methoxy-4-methylphenyl)propanoate (12)

The synthetic procedure for **4** was used in this case. Methyl (*S*)-2-((*tert*-butoxycarbonyl)amino)-3-iodopropanoate (0.2 g, 0.60 mmol) yielded the product **12** (0.110 g, 56% yield).^1^H NMR (500 MHz, CDCl_3_) δ 7.04 (d, *J* = 7.3 Hz, 1H), 6.66 – 6.55 (m, 2H), 4.99 (s, 1H), 4.57 (s, 1H), 3.81 (s, 3H), 3.72 (s, 3H), 3.12 – 2.94 (m, 2H), 2.18 (s, 3H), 1.43 (s, 9H).

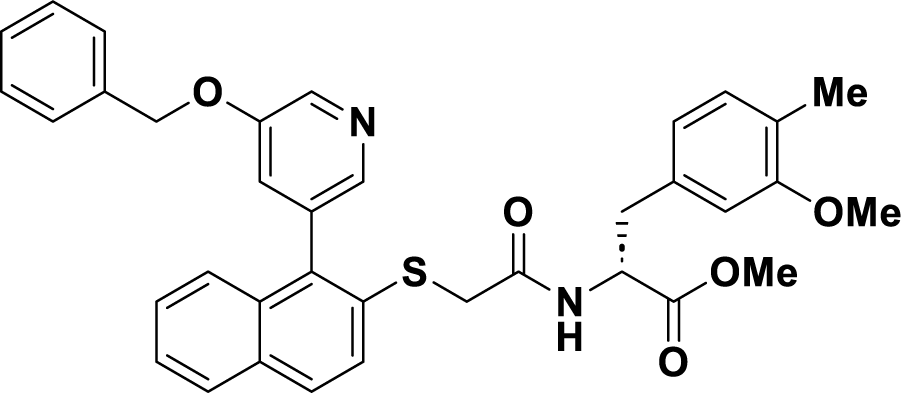

##### Methyl(*R*)-2-(2-((1-(5-(benzyloxy)pyridin-3-yl)naphthalen-2-yl)thio)acetamido)-3-(3-methoxy-4-methylphenyl)propanoate (14)

The synthetic procedure for **10** was used in this case. Ester **9** (0.05 g, 0.11 mmol) yielded product **14** (0.055 g, 78% yield). ^1^H NMR (500 MHz, CDCl_3_) δ 8.43 (dd, *J* = 5.6, 2.8 Hz, 1H), 7.84 – 7.68 (m, 2H), 7.53 – 7.15 (m, 10H), 7.09 – 6.88 (m, 2H), 6.58 – 6.34 (m, 1H), 6.34 – 6.22 (m, 2H), 5.10–01 (m, 1H), 4.93 (q, *J* = 11.6 Hz, 1H), 4.75–4.61 (m, 1H), 3.57 (s, 3H), 3.54–3.51 (m, 2H), 3.49 (s, 3H), 307–2.83 (m, 2H), 1.90 (s, 3H). ^13^C NMR (126 MHz, CDCl_3_) δ 171.5 167.6, 167.5, 157.6, 143.3, 138.1, 138.0, 136.0, 134.3, 134.0, 133.0, 132.8, 131.9, 131.8, 130.5, 129.5, 128.7, 128.0,127.8, 127.7, 127.3, 126.0, 125.5, 123.7, 123.5, 123.0, 120.9, 120.7, 110.4, 70.4, 55.1, 53.5, 37.5, 36.9, 36.7, 15.7.

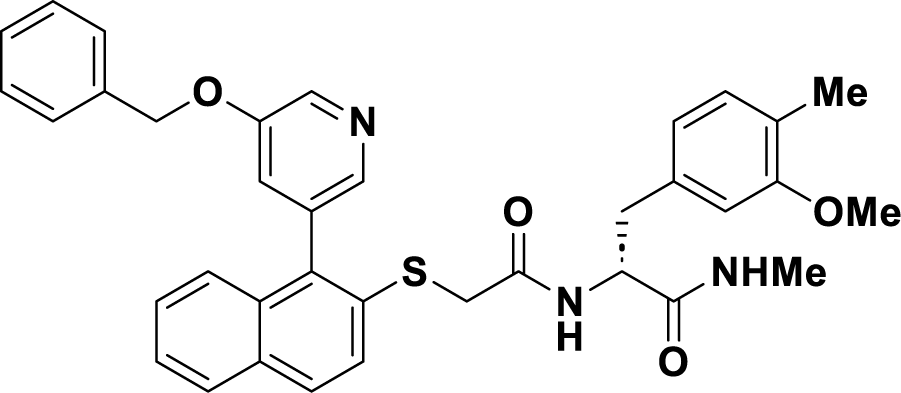

##### (*R*)-2-(2-((1-(5-(benzyloxy)pyridin-3-yl)naphthalen-2-yl)thio)acetamido)-3-(3-methoxy-4-methylphenyl)-*N*-methylpropanamide

The synthetic procedure for **11** was used in this case. Peptide **14** (11 mg, 0.018 mmol) yielded product **15** (8 mg, 80% yield). ^1^H NMR (500 MHz, CDCl_3_) δ 8.53 (s, 1H), 8.16 (s, 1H), 7.94 – 7.72 (m, 2H), 7.50 – 7.28 (m, 10H), 7.13 (s, 1H), 6.86 – 6.72 (m, 1H), 6.63 – 6.39 (m, 2H), 5.46 (d, *J* = 21.9 Hz, 1H), 5.29 – 5.07 (m, 2H), 4.59 – 4.43 (m, 1H), 3.70 (s, 3H), 3.64 (s, 2H), 3.06 – 2.83 (m, 2H), 2.54 (s, 3H), 2.06 (s, 3H). ^13^C NMR (126 MHz, CDCl_3_) δ 170.7, 167.8, 157.8, 154.8, 143.5, 143.3, 138.15, 138.1, 136.0, 134.9, 134.6, 134.3, 132.9, 131.9, 131.7, 130.6, 129.5, 128.7, 128.3, 128.1, 127.7, 127.3, 126.1, 125.4, 124.0, 123.9, 123.5, 120.7, 110.6, 70.5, 55.2, 38.0, 37.19, 37.0, 26.1, 15.8.

HRMS calcd for C36H35N3O4S Na+; 628.2240: HRMS found 628.2243

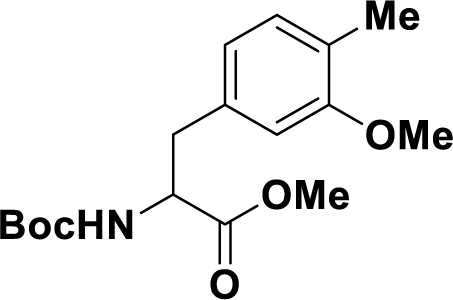

##### Methyl2-((*tert*-butoxycarbonyl)amino)-3-(3-methoxy-4-methylphenyl)propanoate (16)

The synthetic procedure for **4** was used in this case. Methyl 2-((*tert*-butoxycarbonyl)amino)-3-iodopropanoate (0.2 g, 0.60 mmol) yielded the product **16** (0.12 g, 61% yield). ^1^H NMR (500 MHz, CDCl_3_) δ 7.13 – 6.95 (m, 1H), 6.67 – 6.48 (m, 2H), 5.14 – 4.92 (m, 1H), 4.57 (d, *J* = 7.8 Hz, 1H), 3.80 (s, 3H), 3.72 (s, 3H), 3.05 (dd, *J* = 12.9, 6.0 Hz, 2H), 2.18 (s, 3H), 1.42 (s, 9H).^13^C NMR (126 MHz, CDCl_3_) δ 172.4, 157.7, 155.1, 134.6, 130.6, 125.3, 121.0, 110.9, 79.8, 55.2, 54.5, 52.1, 38.2, 28.3(3C), 15.8.

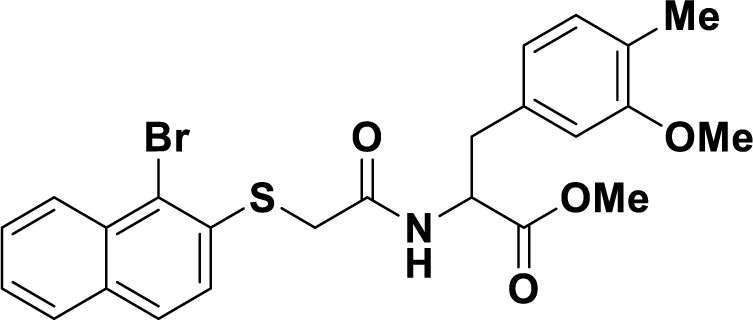

##### Methyl 2-(2-((1-bromonaphthalen-2-yl)thio)acetamido)-3-(3-methoxy-4-methylphenyl) propanoate (18)

The synthetic procedure for **10** was used in this case. Ethyl 2-((1-bromonaphthalen-2-yl)thio)acetate **8** (0.22 g, 0.68 mmol) yielded the coupling product **18** (0.26 g, 76% yield). ^1^H NMR (500 MHz, CDCl_3_) δ 8.23 (dd, *J* = 8.6, 1.0 Hz, 1H), 7.82 – 7.74 (m, 1H), 7.71 – 7.65 (m, 1H), 7.60 (ddd, *J* = 8.4, 6.8, 1.3 Hz, 1H), 7.50 (ddd, *J* = 8.1, 6.9, 1.2 Hz, 1H), 7.18 (dd, *J* = 15.3, 8.4 Hz, 2H), 6.77 – 6.69 (m, 1H), 6.45 (d, *J* = 1.7 Hz, 1H), 6.35 (dd, *J* = 7.5, 1.7 Hz, 1H), 4.81 (ddd, *J* = 7.8, 6.9, 5.4 Hz, 1H), 3.74 (s, 2H), 3.66 (s, 3H), 3.64 (s, 3H), 3.00 (qd, *J* = 14.0, 6.2 Hz, 2H), 2.00 (s, 3H).^13^C NMR (126 MHz, CDCl_3_) δ 171.4, 167.3, 157.7, 134.0, 133.8, 132.5, 132.4, 130.5, 128.4, 128.2, 128.1, 126.6, 126.3, 125.3, 123.7, 121.9, 120.6, 110.3, 55.1, 53.4, 52.2, 37.5, 36.8, 15.7.

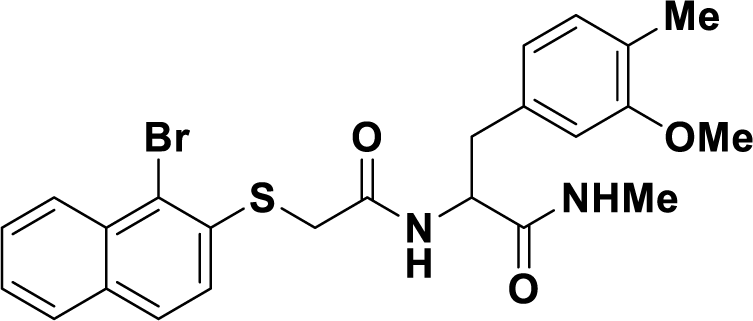

##### 2-(2-((1-bromonaphthalen-2-yl)thio)acetamido)-3-(3-methoxy-4-methylphenyl)-*N*-methylpropanamide (19)

Methyl 2-(2-((1-bromonaphthalen-2-yl)thio)acetamido)-3-(3-methoxy-4-methylphenyl)propanoate (0.250 g, 0.49 mmol) was dissolved in MeOH:CH_2_Cl_2_ (10:1, 11 mL) to it NH_2_Me (40% in H_2_O, 5 mL) was added and reaction was stirred at room temperature for overnight. After completion of reaction the volatile solvents were evaporated off to complete dryness and residue used in next step without purification.

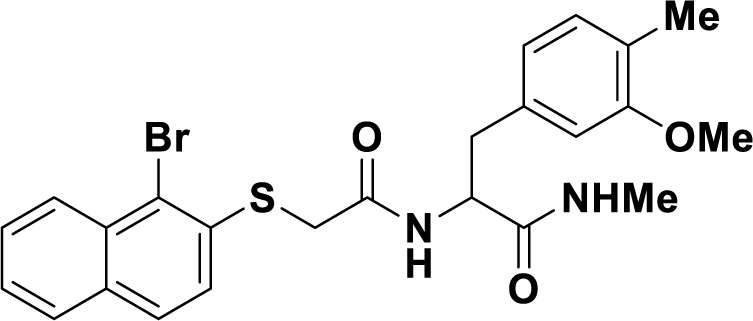

##### 2-(2-((1-(5-(benzyloxy)pyridin-3-yl)naphthalen-2-yl)thio)acetamido)-3-(3-methoxy-4-methylphenyl)-N-methylpropanamide (20)

Using the protocol for compound **9**, above obtained compound yielded the Suzuki coupling product **20** (0.2 g, 67% yield for 2 steps). ^1^H NMR (399 MHz, CDCl_3_) δ 8.51 (t, *J* = 2.7 Hz, 1H), 8.15 (s, 1H), 7.94 – 7.72 (m, 2H), 7.52 – 7.17 (m, 11H), 6.81 – 6.70 (m, 1H), 6.54 (m, 1H), 6.46 – 6.40 (m, 1H), 5.89 – 5.66 (m, 1H), 5.25 – 5.06 (m, 2H), 4.54 – 4.47 (m, 1H), 3.66 (s, 3H), 3.58 – 3.47 (m, 2H), 3.01 – 2.86 (m, 2H), 2.54 (s, 3H), 2.04 (s, 3H). ^13^C NMR (100 MHz, CDCl_3_) δ 170.8, 168.0, 157.8, 154.7, 143.4, 143.2, 138.1,138.0, 136.0, 135.0 134.5, 134.2, 132.9, 131.8, 131.7, 130.5, 129.4, 128.7, 128.63, 128.1,127.7,127.65, 127.3, 126.0, 125.3, 124.01, 123.3, 120.7, 110.6, 70.4, 55.1, 54.9, 38.0, 37.1, 26.0, 15.7. HRMS calcd for C36H35N3O4S Na+; 628.2240: HRMS found 628.224

### GTP turnover assay

The GTP turnover assay was performed using a modified version of the GTPase-GLO^TM^ assay (Promega) as described previously^22,23^. Purified µOR was diluted to 1 µM in 20 mM HEPES pH 7.4, 100 mM NaCl, 0.01% L-MNG, 0.001% CHS, and 20 µM guanosine-5’-triphosphate (GTP) in the presence of various orthosteric (20 µM met-enkephalin, 20 µM naloxone, 20 µM MP, 20 µM H-Tyr-D-Ala-Gly-*N*(Me)Phe-Gly-OH [DAMGO], 20 µM BU72) and allosteric (serially diluted or at excess concentrations of 20 µM for 10105-368-30-605) ligands and incubated for 1.5 hours at room temperature. Concurrently, G_i_ purified in DDM was exchanged by incubating with 1% L-MNG and 0.1% CHS for 1 hour on ice. The exchanged G_i_ was then diluted to 1 µM in 20 mM HEPES pH 7.4, 100 mM NaCl, 0.01% L-MNG, 0.001% CHS, 20 µM guanosine-5’-diphosphate (GDP), 200 µM TCEP, and 20 mM MgCl_2_. Equal volumes of receptor solutions and G_i_ solution were mixed and incubated at room temperature for 60 minutes (agonist-bound receptor experiments) or 90 minutes (apo receptor experiments) with gentle shaking. Controls include mixing equal volumes of both buffers (total initial GTP) and equal volumes of 1 µM G_i_ solution and receptor buffer (intrinsic G protein turnover). Equal volume of GTPase-Glo reagent supplemented with 10 µM adenosine 5’-diphosphate (ADP) in 20 mM HEPES pH 7.4, 100 mM NaCl, 0.01% MNG and 0.001% CHS was added and incubated with gentle shaking for 30 minutes, followed by addition of further equal volume of detection reagent. After brief (10 minute) incubation, luminescence was measured with a MicroBeta counter.

#### *In-cell* cAMP assay

HEK293 cells stably expressing the Epac1-based FRET cAMP biosensor (PMID: 23374187) were transfected with a pcDNA3.1(+)neo-based plasmid encoding a FLAG-tagged µOR using Fugene6 (Promega E2691) according to the manufacturer’s instructions. Cells were maintained in growth medium [D-MEM, Gibco 10566016; supplemented with 10% fetal bovine serum, Gibco 10270106; 1% sodium pyruvate, Gibco 11360039; 1% MEM non-essential amino acids, Gibco 11140068; and 1% penicillin– streptomycin Solution, Gibco 15140122] supplemented with 50 µg/ml zeocin (for selection of the Epac1-based FRET cAMP biosensor) and 500 µg/ml G418 (for selection of FLAG-tagged µOR). Following limited dilution cloning in growth medium containing zeocin and G418, single colonies were picked and tested for their ability to decrease forskolin-induced cAMP formation upon activation with met-enkephalin (as described below). One clone was chosen, propagated and used for subsequent testing.

The day before assaying cells were seeded at 100 µL per well in a black poly-L-lysine coated 96-well plate at 30,000 cells per well. Prior to assaying, cells were washed with 100 µL HBSS buffer (HBSS Gibco 14025, 20 mM HEPES pH 7.5, 1 mM CaCl2, 1 mM MgCl2, 0.1% BSA), and replaced with 25 µL HBSS buffer in each well. For testing of the allosteric modulator ability to affect agonist-induced cAMP formation, the allosteric modulators were added to wells (25 µl, 6x final concentration) and preincubated for 20 min. Forskolin, diluted in a volume of 50 µl, was then added to wells to final a concentration of 15 µM and incubated for 20 min while following the change in FRET ratio between CFP and YFP as a measure of cAMP formation using an Envision plate reader (PMID: 23374187). Then orthosteric agonists were diluted in forskolin buffer (corresponding to a final concentration of 15 µM), added at a volume of 50 µl to wells, and the decrease in cAMP formation followed for 1 hr. For quantification of cAMP levels, the AUC for each well was calculated. For NAM testing, a concentration of met-enkephalin or morphine corresponding to EC_80_ was added, and the NAM potency (IC50) determined by non-linear regression after fitting the AUC from the point of agonist addition as a function of the NAM concentration to a four-parameter logistic curve using GraphPad Prism. For determining the effect of allosteric modulator alone, then orthosteric agonist addition step was omitted, and the AUCs from the addition of forskolin alone was quantified.

### TRUPATH Assays

The allosteric activity for the highly enriched μOR NAM (Compound **368**) was tested in combination with various orthosteric agonists (morphine, fentanyl, and met-enkephalin) or antagonist (naloxone). To determine Compound **368**’s negative allosteric effect on the coupling of the μOR and the heterotrimeric G proteins, HEK293T cells were co-transfected in a 5:1:5:5 ratio of μOR, individual Gα-RLuc8 (G_i1_ or G_i3_), Gβ_3_, and GFP2-Gγ_9_ construct, while a combination of a 5:1:5:5 ratio of μOR, individual Gα-RLuc8 (G_i2_, G_oA_, or G_oB_), Gβ_3_, and GFP2-Gγ_8_ construct and a 5:1:5:5 ratio of μOR, individual Gα-RLuc8 (G_z_), Gβ_3_, and GFP2-Gγ_1_ construct was used in the presence of transfection reagent, Transit 2020. The next day, the transfected cells were plated and incubated overnight into poly-L-lysine coated 96-well white clear bottom cell culture plates at a density of 50,000 cells per 200 μl per well using Dulbecco’s modified Eagle’s medium supplemented with 1% dialyzed fetal bovine serum. On the day after, cells were washed with 60 μL of a drug buffer (1× HBSS and 20 mM HEPES, pH 7.4) per well after aspirating the cells media. To measure the allosteric activity of Compound **368** with individual orthosteric agonists, 60 μL of 7.5 μM RLuc substrate coelenterazine 400a (Nanolight) was added per well, followed by the addition of 30 μL per well of 3X final concentrations (0 and 90 μM) of Compound **368** that was prepared in a drug buffer supplemented with 0.3% BSA and incubated for 20 min in the dark at room temperature. Finally, 30 μL of 4X final various concentrations of morphine, fentanyl, or met-enkephalin were prepared in a drug buffer containing 0.3% BSA then added and incubated for 5 min. To measure the allosteric activity of Compound **368** with naloxone, 30 μL of 2X final various concentrations of naloxone was added per well, followed by the addition of 30 μL per well of 2X final concentrations (0 and 90 μM) of Compound **368** that was prepared in a drug buffer supplemented with 0.3% BSA and incubated for 20 min. Afterward, 60 μL of 7.5 μM RLuc substrate coelenterazine 400a (Nanolight) was loaded per well and incubated in the dark at room temperature. After 5 min, 30 μL of 5X final concentration of DAMGO’s EC_80_ prepared in a drug buffer that contained 0.3% BSA was loaded per well and incubated in the dark at room temperature for an additional 5 min. Subsequently, Mithras LB940 multimode microplate reader was used to measure the BRET ratios for Gα protein activation by quantifying the ratio of the GFP2 emission to RLuc8 emission for 1s per well at 510 nm and 395 nm, respectively. GraphPad Prism 9 software was used to determine the potency and efficacy of the examined orthosteric agonists and the antagonist by plotting their different used concentrations against the normalized BRET ratios that were normalized by defining 0% as basal and 100% as maximum values at 0 μM **368**.

### Formation & purification of µOR- κ_ICL3_/naloxone/Nb6/10105-368-30-605 complex for cryo-EM studies

µOR- κ_ICL3_ was diluted to 20 µM in 20 mM HEPES pH 7.5, 100 mM NaCl, 0.01% L-MNG, 0.003% GDN, 0.0013% CHS, 200 µM naloxone and 200 µM 10105-368-30-605 and incubated for ∼30 minutes followed by addition of 2:1 molar excess of Nb6 and further incubation on ice overnight. Excess Nb6 was removed by M1 anti-FLAG immunoaffinity chromatography, and the pure µOR- κ_ICL3_/Nb6/naloxone/10105-368-30-605 complex was eluted in a buffer composed of 20 mM HEPES pH 7.5, 100 mM NaCl, 0.00075% L-MNG, 0.00025% GDN, 0.0001% CHS, 100 µM naloxone, 100 µM 10105-368-30-605, FLAG peptides and 5 mM EDTA. Complex was concentrated to ∼16 mg mL^-1^ for electron microscopy and analysis by SDS-PAGE.

### Cryo-EM data acquisition & processing

#### µOR- κ_ICL3_/naloxone/Nb6/10105-368-30-605

The µOR- *κ_ICL3_*/naloxone/Nb6/10105-368-30-605 complex prepared above was applied to glow-discharged 200 mesh grids (Quantifoil R1.2/1.3) at a concentration of ∼16 mg mL^-1^ and vitrified using a Vitrobot Mark IV (Thermo Fisher Scientific) at 100% humidity at 22 °C after blotting for 3 s with a blot force of 3. CryoEM images were collected on a Titan Krios (Thermo Fisher Scientific) equipped with X-FEG and Selectris X image filter operated at 300 kV at a nominal magnification of 165,000x using a Falcon 4 camera in counting mode corresponding to a pixel size of 0.741 Å. A total of 10,007 image stacks were obtained with a dose rate of 5.0 e^-^/pixel/s and total exposure time of 6.6 s, resulting in a total dose of 60 electrons per Å^2^. The defocus range was set to -0.6 to -1.4 µm.

Dose-fractionated image stacks were imported to Relion^51^ and subjected to patch-based beam-induced motion correction using MotionCor2^52^ and CTF parameters were determined with CTFFIND4^53^. After removing micrographs with an estimated CTF resolution of less than 3.4 Å, a total of 4,254,557 particles were picked using a template-based auto-picking protocol. These particles were imported to CryoSPARC^54^ for several rounds of 2D classification, resulting in 898,414 particles. These particles were then subjected to iterative rounds of 3D classification using *ab initio* classification with concurrent 2D classification rounds to “rescue” particles from “bad” 3D classes. After a final round of 2D classification, a set of 128,613 final particles were reconstructed to 3.26 Å nominal resolution at FSC value of 0.143 with local refinement. Local resolution was estimated with CryoSPARC.^54^

#### Model building & refinement

The initial template for the µOR- κ_ICL3_/naloxone/Nb6/10105-368-30-605 structure was derived from the µOR- κ_ICL3_/alvimopan/Nb6 structure solved previously.^28^ The initial model was docked into the EM density map using UCSF Chimera followed by iterative model building in Coot.^55^ Ligand coordinates (naloxone, 10105-368-30-605) and geometry restraints were initially generated using phenix.elbow.^56^ The final model was subjected to global refinement and minimization in real space with phenix.real_space_refine.^57^ Residues with weak side chain density were stubbed while preserving sequence information. Molprobity^58^ was used to evaluate model geometry. The final pose of the allosteric modulators for each structure was determined using a docking into density protocol implemented in Glide (Schrödinger) followed by evaluation with all-atom molecular dynamics simulations (described below). The final refinement parameters are provided in Table S5 for µOR- κ_ICL3_/naloxone/Nb6/10105-368-30-605.

### MD simulations

#### System setup

Simulation systems were built with CHARMM-GUI^59^ using the Membrane Builder^60–63^. The chains for the receptor and ligands from structure presented here (naloxone and Cmpd **368** for the µOR- κ_ICL3_/naloxone/Nb6/10105-368-30-605) were selected and other proteins (e.g. Nb6) were removed. The CHARMM General Force Field^64,65^ was used to generate topology and parameter files for the small molecule ligands (Cmpd **368** and naloxone). Missing side chains were modeled in the CHARMM-GUI and the receptor chains were capped with neutral N-terminal acetyl groups and C-terminal methylamide groups. The N- and C-terminus of met-enkephalin were patched to standard N- and C-termini. Titratable groups were simulated in their dominant protonation state for pH 7. The receptors were placed in the membrane by aligning the first principal axis of the receptor to the z-axis, followed by visual inspection. The aligned structures were then inserted into a palmitoyl-oleoyl-phosphatidylcholine (POPC) bilayer with a minimum water height on either side of the bilayer of 22.5 Å and a bilayer area chosen to be ∼4 times that of the protein surface, normalizing the upper and lower leaflet surface area by additional POPC molecules accordingly. 150 mM potassium chloride (KCl) ions were added to neutralize the system charge.

#### Simulation protocol

All simulations were performed with the CHARMM36m force field^66^ for proteins, ions and lipids, and the TIP3 model^67^ was used for waters. Simulations were performed using OpenMM^68^ on single graphics processing units (GPUs). The system was equilibrated initially with 1 fs time steps (125 ps per run for 3 runs) followed by 2 fs time steps (500 ps per run for 3 runs) for a total of 6 equilibration steps each with continually decreasing force constants applied to the protein backbone (10.0, 5.0, 2.5, 1.0, 0.5, 0.1 kcal/mol Å^2^), side chains (5.0, 2.5, 1.0, 0.5, 0.1, 0.0 kcal/mol Å^2^), waters and lipids (2.5, 2.5, 1.0, 0.5, 0.1, 0.0 kcal/mol Å^2^) and ions (10.0, 0.0, 0.0, 0.0, 0.0, 0.0 kcal mol Å^2^). Production runs were performed in NPT ensemble at 310.15 °K at 1 bar with 4 fs time steps employing hydrogen mass repartitioning (HMR)^69^. Bonds to hydrogen atoms were constrained with the SHAKE algorithm^70^. A switching distance of 10 Å and cutoff distance of 12 Å were used for nonbonded interactions, and long-range electrostatics were computed with particle mesh Ewald (PME). Coordinates were written out every 100 ps.

#### *In vivo* experiments

All experiments used male C57BL/6J mice (Jackson Laboratories Bar Harbor) (20-35 g) at 5 per cage with a 12-hour light/dark cycle (i.e. lights off at 7:00 PM and on at 7:00 AM) with *ad libitum* access to food and water. Mice were kept at a constant temperature (20-24 °C) and relative humidity (40-50%). All studies reported here are in compliance with ARRIVE guidelines^71^ and the National Institutes of Health guide for care and use of animals (NIH Publications No. 8023, revised 1978) and were approved and conducted in agreement with the Institutional Animal Care and Use Committees at the University of Florida. Mice were used in pharmacokinetics^11^, warm-water tail withdrawal^72,73^, locomotor and respiration (via comprehensive lab animal monitoring system, CLAMS)^72,73^, conditioned place preference^72,74^, and naloxone-precipitated opioid withdrawal assays^38^, all described in detail below.

#### Compound treatment

Compound **368** (racemic mixture) was synthesized as above. Other drugs (morphine, fentanyl, and naloxone) were purchased and used as previously described^11,38^.

#### Pharmacokinetics of Cmpd 368

Cmpd **368** was administered to mice intravenously at 10 mg/kg dose in four C57BL/6J mice for each time point. After 30, 60, or 120 minutes, mice were anesthetized with isoflurane followed by removal of blood and the animals were sacrificed for removal of the brain. Brains were rinsed with PBS, dried, flash-frozen and weighed for subsequent studies on brain penetration of Cmpd **368**. Tissue samples were placed into Navy bead lysis kit tubes and compared against naïve tissue, which was used for standard, quality control and blank samples. An appropriate volume of cold acetonitrile:water (3:1) was added to each sample tube to normalize the concentrations to 250 mg/mL. Tubes were then placed into a bead heater for 3 minutes, followed by centrifugation at 3200 rpm for 5 minutes at 4 °C. The resulting supernatants were transferred to Eppendorf tubes and stored at -80 °C until analysis.

For analysis, samples were thawed on ice and extensively mixed, followed by further centrifugation at 3200 rpm for 5 minutes at 4 °C. 30 µL of supernatant was collected and transferred to a 96 well plate along with 30 µL standards, quality controls, blanks, and double blanks. Cold acetonitrile (150 µL) spiked with internal standard was then added to blanks, standards, quality controls and unknown samples, while cold acetonitrile alone was added to double blanks. Samples were mixed for 10 minutes followed by further centrifugation at 3200 rpm for 10 minutes at 4 °C. Supernatants after centrifugation were evaporated to dryness in a 96 well plate, followed by reconstitution with 100 µL 0.1% v/v formic acid in water:acetonitrile (90:10). The plate was sealed, vortexed, and briefly centrifuged to settle the samples, and was submitted for LC/MS analysis as previously described^11,36^ using a mass spectrometer (SCIEX Triple Quad 5500+ system – QTRAP ready) linked to a liquid chromatography system (ExionLC AD – HPLC).

#### Antinociceptive testing

The 55 °C warm-water tail-withdrawal assay was performed as previously described^37,75^. Latency of the mouse to remove its tail from warm water was recorded as the endpoint. After determining baseline tail-withdrawal latencies, mice were administered opioid agonists with or without subcutaneous pretreatment with vehicle (10% DMSO/10% Solutol/80% saline, 0.9%), a graded dose of Cmpd **368** (1-100 mg/kg) and/or naloxone. Vehicle or Cmpd **368** was administered 30 minutes before administering saline or naloxone (0.1, 1 or 10 mg/kg) 15 min in advance of morphine (10 mg/kg, s.c), at which point the tail-withdrawal assay was performed every 10 minutes until a return to baseline responses were achieved (out to 150 minutes). To avoid excess tissue damage, mice that failed to withdraw their tails after 15 seconds had their tails removed from the bath and were assigned a maximum antinociceptive response (100%, see below equation). The assay was additionally performed with fentanyl as the orthosteric opioid agonist, but fentanyl was administered at 0.4 mg/kg. At each time point, antinociception was then calculated as follows:

> % antinociception = 100 * ((test latency – baseline latency)/(15 – baseline latency))

#### Locomotor and respiration effects

Locomotor activity (as ambulations) and respiratory depression (as breaths per minute) were measured using automated, closed-air Comprehensive Lab Animal Monitoring system (CLAMS) in a similar manner to that previously described^37,38,76^. Briefly, mice were habituated in chambers for 60 minutes, during which baselines for each animal were measured. Vehicle or increasing doses of Cmpd **368** (1-100 mg/kg) was then administered, followed 30 min later by administration of either vehicle, morphine (10 mg/kg), or morphine with low-dose naloxone (0.1 mg/kg), and confined 5 min later in individual CLAMS testing chambers for measures lasting 200 minutes. The respiration rate (breaths per minute) was measured via pressure transducers built into the sealed CLAMS cages, whereas infrared beams located in the cage floors measured ambulations via sequential beam breaks. Data are expressed as a percentage of vehicle responses ± SEM for ambulations or breaths per minute, averaged over 20-min periods for 140 min post-injection of the test compound.

#### Conditioned place preference (CPP) assay

Groups of mice were place conditioned using a counterbalanced design in a three-chamber place conditioning system (San Diego Instruments) as described previously^36,72^. The amount of time individual mice spent in each compartment both before and after place conditioning was measured over a 30-min testing period, where animals freely explored the three compartments. Following determination of initial preference, mice were administered 0.9% saline, and half the sample consistently confined in a randomly assigned outer chamber for 40 min. Four hours later, mice were pretreated subcutaneously for 30 min with either vehicle or Cmpd **368** (100 mg/kg), followed by all mice receiving low-dose naloxone (0.1 mg/kg, s.c.). Fifteen minutes later, mice were administered morphine (10 mg/kg, i.p.) and place conditioned for 40 minutes in the opposite outer chamber. Mice were place conditioned in this manner for 2 days, with a final place preference performed on the fourth day. Data are plotted as the difference in time spent in the eventual morphine-paired chamber versus the vehicle-paired compartment.

#### Naloxone-precipitated opioid withdrawal assay

Mice were made physically dependent on morphine using previously established methods^37,38^. Briefly, mice were placed into four groups (n=10-11 per group) and were given either saline (group **1**) or escalating doses of morphine (10-75 mg/kg, groups **2-4**) twice daily for four days. A final dose of saline (for group **1**) or morphine (25 mg/kg) was given on day 5 to assess antinociceptive tolerance. Two hours later, precipitated withdrawal symptoms were assessed for 15 min following various treatments. Groups **1** and **2** were given a conventional dose of naloxone (10 mg/kg), while groups **3** and **4** were pretreated 30 min with either saline (**3**) or Cmpd **368** (**4**, 100 mg/kg, s.c.) ahead of administration of low-dose naloxone (0.1 mg/kg). Immediately after the various naloxone treatments, withdrawal behaviors were quantified from mice in 16 x 45 cm plexiglass cylinders for 15 minutes following established protocols^37,38^.

### Figure Preparation

Protein structure figures were created using the *PyMOL* Molecular Graphics System Version 2.3.5 Schrödinger (http://pymol.org). Density map figures were created using UCSF *Chimera*^77^ and UCSF *Chimera X*^78^. Graphs were created using *GraphPad Prism.* Schematic figures were created using *BioRender*. Molecular dynamics trajectories were analyzed with *MDAnalysis*^79,80^. Figures were constructed in *Adobe Illustrator*.

## Supplementary Figures

**Figure S1.**
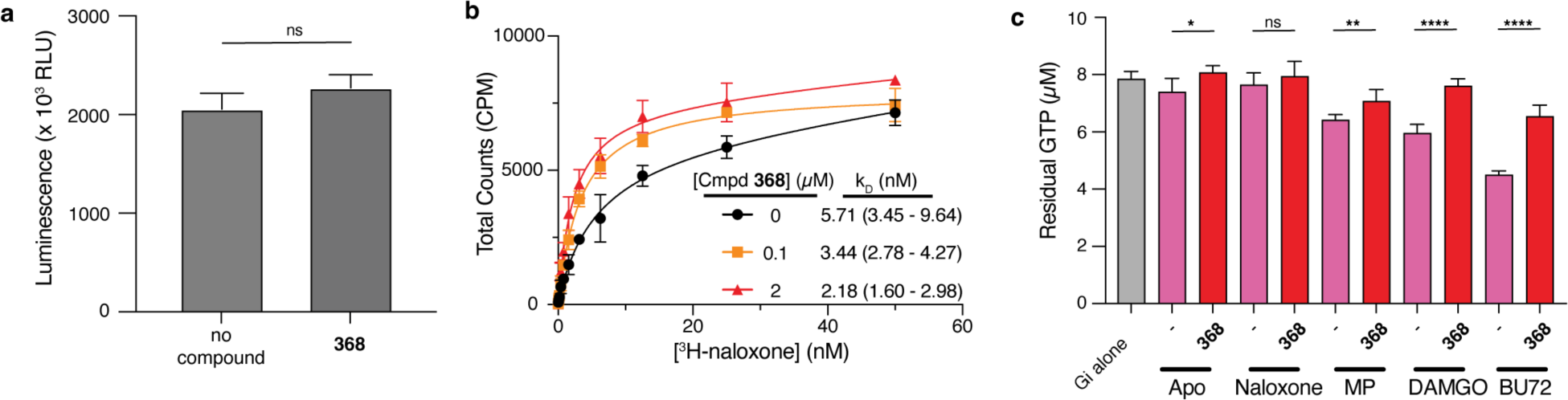
Initial biochemical characterization of µOR allosteric modulators from DEL screen. (a) Excess concentrations of 368 have no impact (P > 0.05, unpaired t-test) on Gi1 intrinsic turnover in the absence of receptor. Error bars correspond to standard deviations of four measurements. (b) We show, using a direct 3H-naloxone binding experiment, that increasing concentrations of 368 result in an increased antagonist affinity for µOR-containing membranes. Fitted affinity values are shown along with 95% confidence intervals. Error bars represent the standard deviations of four measurements (c) The GTP turnover assay was used to show that 20 µM 368 inhibits turnover for a wide variety of orthosteric site conditions, ranging from weak (naloxone) to moderate (mitragynine pseudoindoxyl, MP) partial agonists to peptide (DAMGO) or small molecule (BU72) full agonists (all orthosteric molecules also present at 20 µM). P values are denoted as follows: ns (P>0.05), * (P≤0.05), ** (P≤0.01), *** (P≤0.001), and **** (P≤0.0001) and were calculated using an unpaired t test assuming Gaussian distributions.

**Figure S2.**
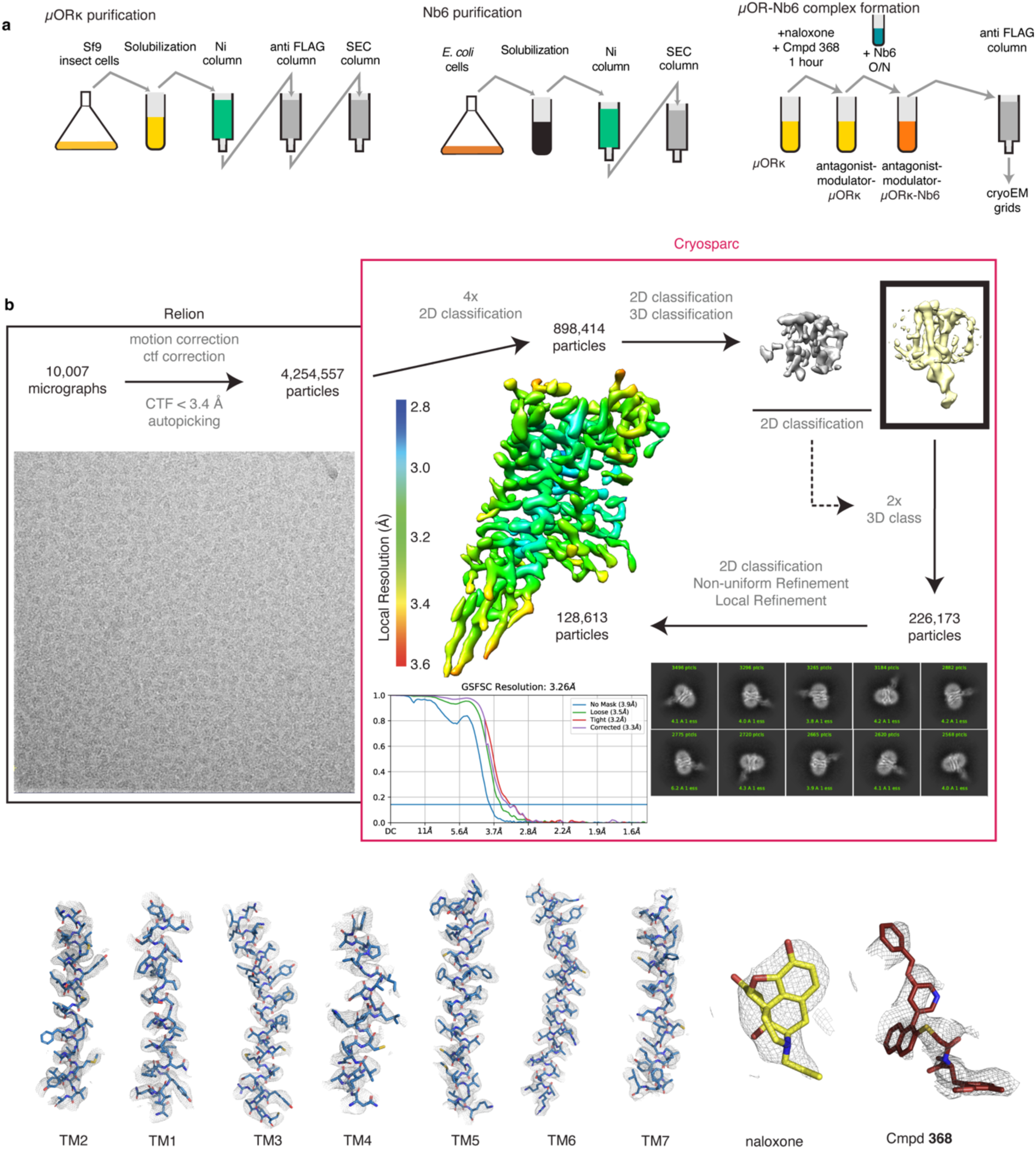
CryoEM structure determination of µORκ-Nb6 bound to naloxone and Cmpd *368.* (**a**) Schematic of µORκ purification and complex formation with Nb6. (**b**) Cryo-EM data collection and processing pipeline, showing representative micrographs of the µORκ-Nb6 complex, reference-free 2D cryo-EM class averages, and processing flow chart. This includes motion and CTF correction in Relion, followed by particle selection, 2D and 3D classifications, density map reconstructions, “gold standard” FSC curves in Cryosparc, and the final density map colored by local resolution. Also included is the cryo-EM density map and model for the seven transmembrane helices of the µORκ, as well as naloxone and **368**.

**Figure S3.**
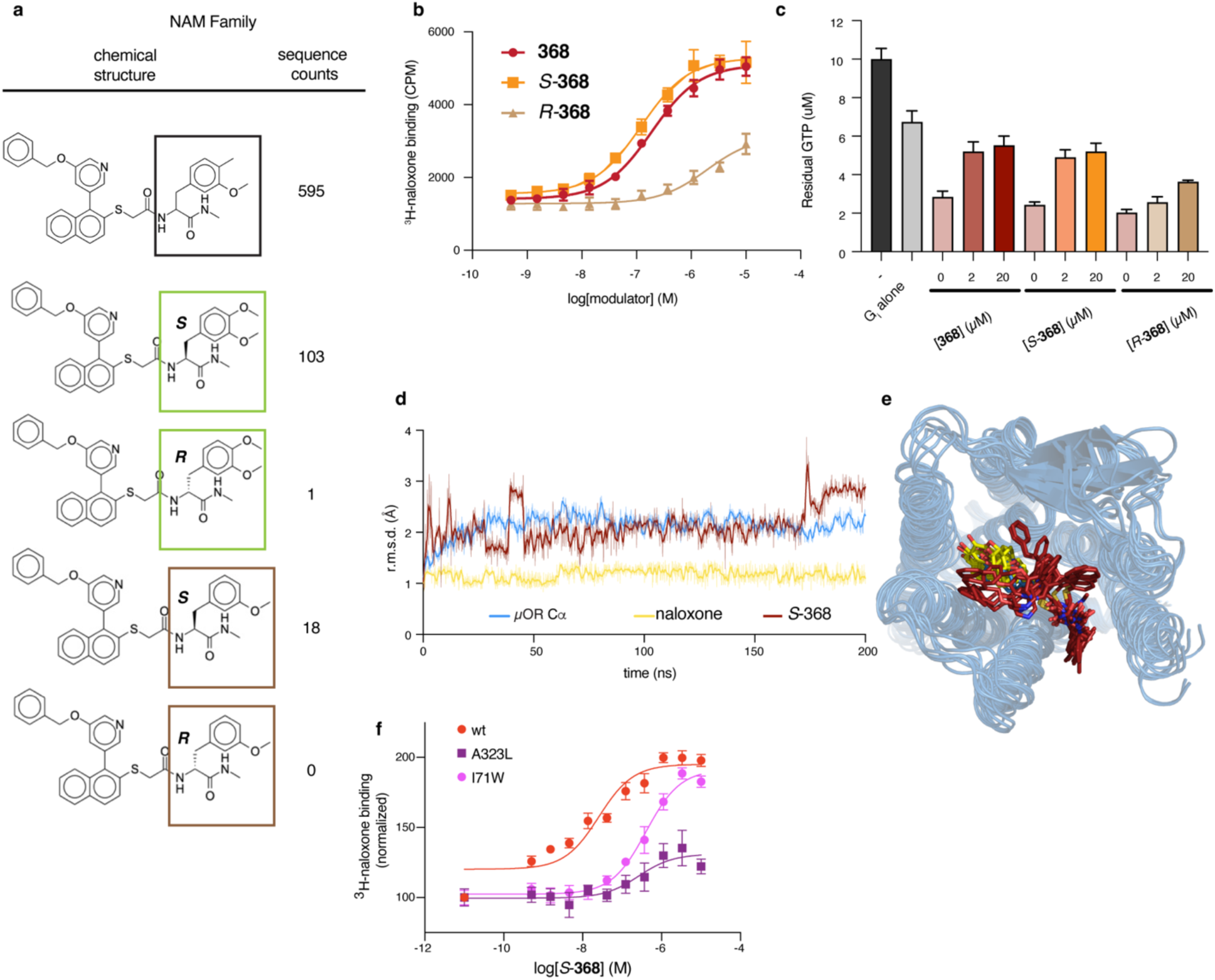
Stereochemistry & MD analysis of *368*. (**a**) Comparison of the chemical structures and raw sequencing counts in the DEL selections for a series of molecules in the enriched **368** family. *S*-stereoisomers are substantially more enriched than their *R*-counterparts. (**b**) Comparison of the individually-synthesized stereoisomers of the racemic **368** “hit”, demonstrating that the *R*-**368** is >100 fold weaker than the S-**368** isomer, consistent with the DEL enrichment data. (**c**) GTP turnover assay again demonstrating that *S*-**368** retains the ability to potently inhibit fentanyl (5 µM)-induced GTP turnover like the racemic **368** “hit”, while the *R*-**368** isomer does not display full inhibition even at 20 µM. Accordingly, the S-isomer of 386 was modeled and docked into the cryo-EM density map which was then subjected to MD simulation for 200 ns (without Nb6). (**d**) The overall root mean square deviations (rmsd) throughout the trajectory were calculated for protein Cα (blue) and all atoms in **368** (red) and naloxone (yellow). **368** remains in broadly the same conformation for the duration of the simulation as observed for 10 ns simulation snapshots (**e**) with low rmsd, though some heterogeneity is observed for the phenyl moiety of the NAM. Naloxone is very rigid throughout the trajectory. (**f**) Titration of S-**368** against either wild type µOR (red), A323L µOR (purple) or I71W µOR (pink) expressing membranes demonstrates that both mutations substantially inhibit the ability of the NAM to enhance ^3^H-naloxone affinity.

**Figure S4.**
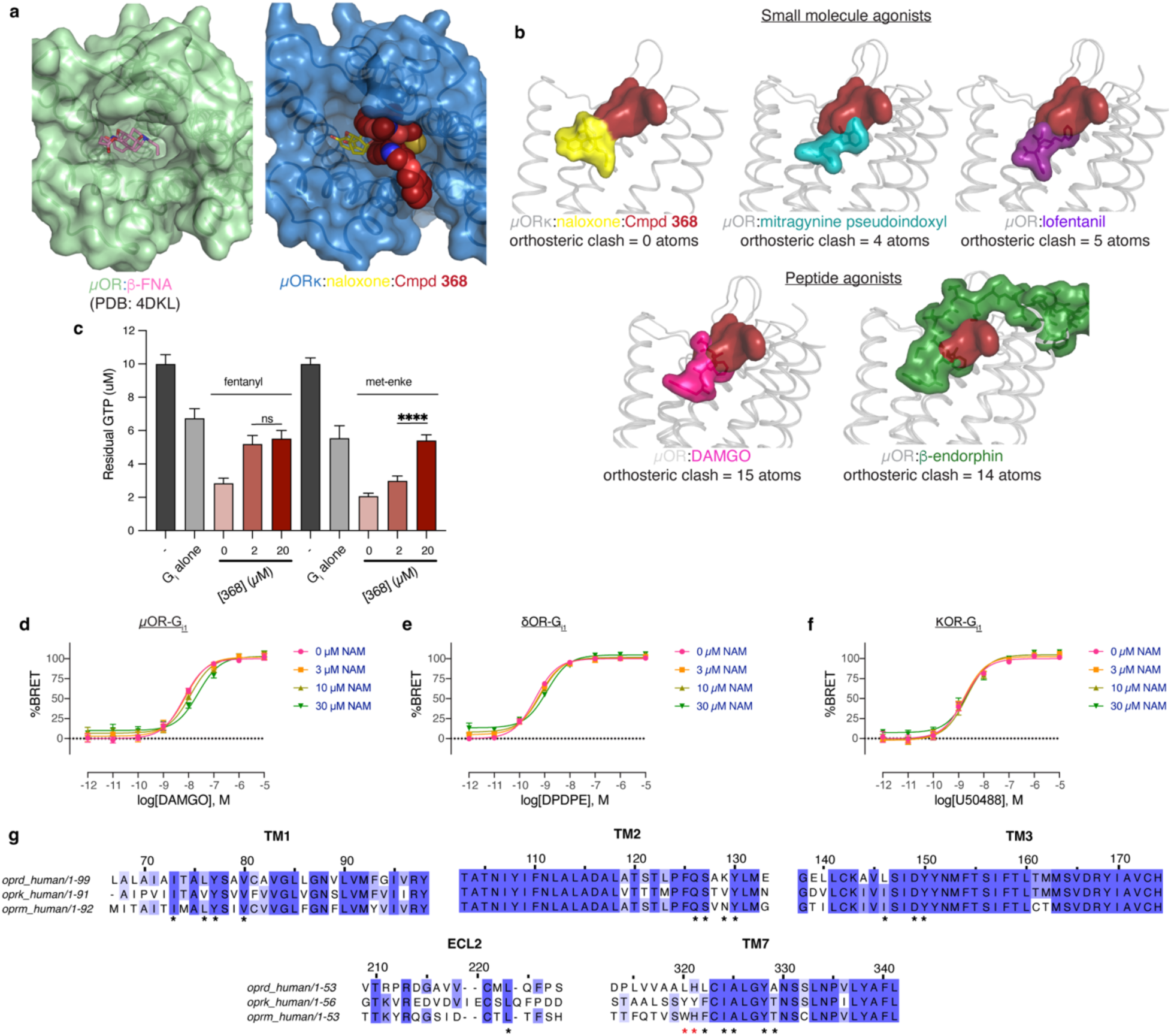
Probe dependence of *368*. (**a**) Comparison of the extracellular vestibule regions of the previous β-FNA-bound µOR and the current naloxone-NAM-bound µOR. The presence of **368** sterically restricts the ability of small molecule antagonists to enter/exit the orthosteric site. (**b**) Alignment of the receptor regions of various ligand-bound µOR structures demonstrate that small molecule orthosteric compounds (top; naloxone [present work], lofentanil [PDB: 7T2H], MP [PDB:7T2G]) have little or no steric clash with **368** (calculated as the number of atoms in the orthosteric ligand within 1 Å of **368**), while both peptide agonists (bottom; β-endorphin [PDB: 8F7Q], DAMGO [PDB: 6DDE]) have clear and substantial steric clash when overlaid with the **368** binding site. (**c**) Additional **368** past 2 µM does not result in increased inhibition of fentanyl-induced GTP turnover but does further inhibit met-enkephalin-induced turnover, perhaps because **368** binds allosterically with small molecules and competitively with peptide agonists. Orthosteric agonists present at 5 µM. (**d-f**) TRUPATH assays for G_i1_ activation for three different opioid receptor/ligand pairs; (**d**) µOR/DAMGO, (**e**) *δ*OR/DPDPE, (**f**) κOR/U50488. Fit parameters at different NAM concentrations are shown below the curves. **368** has the largest impact on DAMGO activation of µOR and shows some activity against *δ*OR activation by DPDPE, but has no effects on κOR activation by U50488. (**g**) Sequence alignment of human µOR, *δ*OR and κOR at structural elements with interaction with **368** (residues within 6 Å denoted with an *). Numbering refers to positions within the human µOR. Red asterisks denote residues with important interactions with **368** but have side chains that are predicted to clash or no longer form productive interactions with **368**. P values are denoted as follows: ns (P>0.05), * (P≤0.05), ** (P≤0.01), *** (P≤0.001), and **** (P≤0.0001) and were calculated using an unpaired t test assuming Gaussian distributions.

**Figure S5.**
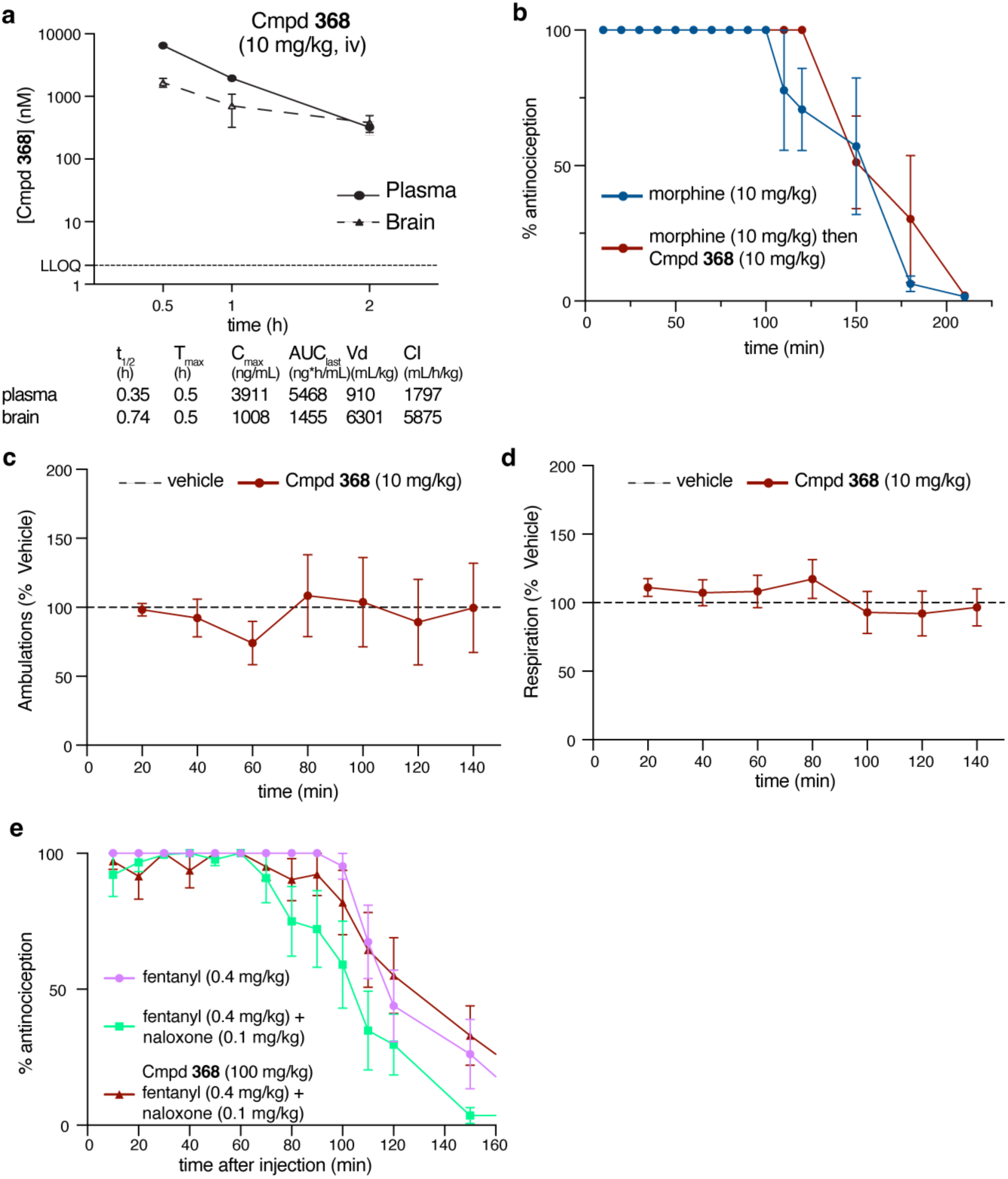
*In vivo* behavior of 368. (**a**) Pharmacokinetics measurements of **368** at 10 mg/kg administered intravenously. **368** enters the brain with a maximum concentration of 1.66 uM, well past the observed affinity of the compound from radioligand binding (133 nM), but significantly lower than the concentration needed to inhibit agonist activity in cell assays. (**b**) In the absence of naloxone, **368** does not have significant effects on morphine-induced antinociception in the 55 °C warm-water tail-withdrawal assay (*F*_(14,84)_=0.80, p=0.67; Two-way RM ANOVA). This contrasts to the observed potentiated antagonism when in the presence of a low-dose (0.1 mg/kg) of naloxone (Fig. 5b). The CLAMS assay demonstrates that in the absence of orthosteric compounds, **368** alone (10 mg/kg) has no significant impact on ambulation (*F*_(6,108)_=0.36, p=0.90; Two-way RM ANOVA) (**c**) and respiration (*F*_(6,108)_=1.19, p=0.32; Two-way RM ANOVA) (**d**). (**e**) While low-dose naloxone (0.1 mg/kg) treatment has some ability to inhibit fentanyl induced antinociception from 110-150 min post administration in the tail-flick assay (pink vs. green lines; *F*_(24,318)_=2.03, p=0.003; Two-way RM ANOVA w/Tukey post hoc test), pre-administration of **368** (100 mg/kg, s.c.) proved unable to potentiate low-dose naloxone antagonism to further inhibit fentanyl analgesia (red vs pink line; *F*_(13,182)_=0.87, p=0.58; Two-way RM ANOVA with Sidak post hoc test), in contrast to morphine (Fig. 5b).

## Supplementary Tables

**Table S1.**
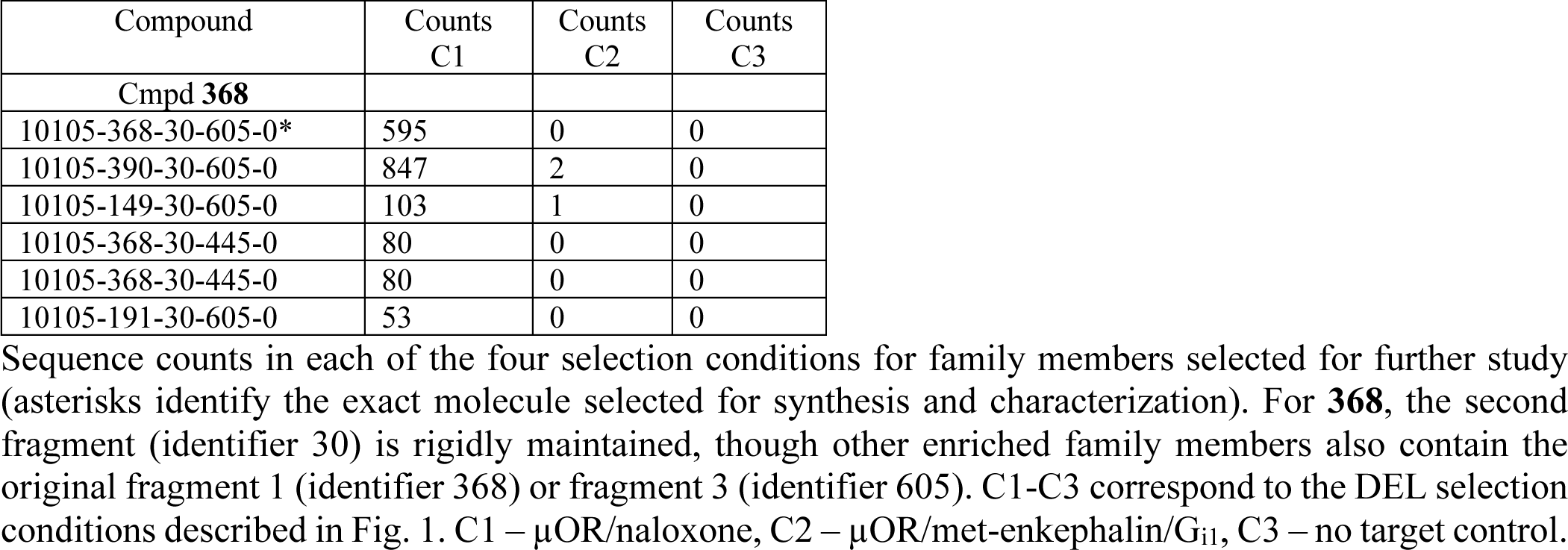
Enrichment properties of selected 368 family members.

| Compound | Counts<br>C1 | Counts<br>C2 | Counts<br>C3 |
| --- | --- | --- | --- |
| **Cmpd 368** |  |  |  |
| 10105-368-30-605-0* | 595 | 0 | 0 |
| 10105-390-30-605-0 | 847 | 2 | 0 |
| 10105-149-30-605-0 | 103 | 1 | 0 |
| 10105-368-30-445-0 | 80 | 0 | 0 |
| 10105-368-30-445-0 | 80 | 0 | 0 |
| 10105-191-30-605-0 | 53 | 0 | 0 |

**Table S2.** Chemical properties of 368.

| MW | logP | PSA | ROTN | HBD | HBA | RCount | ARCount |
| --- | --- | --- | --- | --- | --- | --- | --- |
| 605.76 | 6.36 | 89.55 | 12 | 2 | 7 | 5 | 5 |

MW; molecular weight (Da). logP; predicted octanol/water partition coefficient. PSA; polar surface area ( $\text{\AA}^2$ ). ROTN; rotatable bonds. HBD; hydrogen bond donors. HBA; hydrogen bond acceptors. RCount; number of rings. ARCount; number of aromatics.

**Table S3:** Summary of TRUPATH naloxone potency values.

| Ligand | **368** (μM) | pEC<sub>50</sub> ± SEM (BRET-Gi1 protein activation), M |
| --- | --- | --- |
| Naloxone | 0 | 7.82 ± 0.21 |
| | 90 | 8.69 ± 0.39 |

**Table S4:** Summary of TRUPATH potency values for G_i/o/z_ activation.

| pEC<sub>50</sub> ± SEM, M | **368** (μM) | Orthosteric μOR agonists |  |  |
| --- | --- | --- | --- | --- |
|  |  | Morphine | Fentanyl | Met-enkephalin |
| Gαi1 | 0 | 8.29 ± 0.15 | 8.57 ± 0.11 | 8.64 ± 0.10 |
|  | 90 | 7.09 ± 0.17 | 7.31 ± 0.10 | 7.44 ± 0.14 |
| Gαi2 | 0 | 8.01 ± 0.22 | 8.65 ± 0.17 | 8.61 ± 0.16 |
|  | 90 | 6.95 ± 0.26 | 7.33 ± 0.23 | 7.26 ± 0.32 |
| Gαi3 | 0 | 8.06 ± 0.18 | 8.43 ± 0.15 | 8.43 ± 0.14 |
|  | 90 | 6.72 ± 0.25 | 7.15 ± 0.12 | 7.28 ± 0.10 |
| GαoA | 0 | 8.40 ± 0.15 | 8.83 ± 0.11 | 8.82 ± 0.10 |
|  | 90 | 7.28 ± 0.17 | 7.71 ± 0.19 | 7.60 ± 0.22 |
| GαoB | 0 | 8.47 ± 0.11 | 8.75 ± 0.10 | 8.89 ± 0.11 |
|  | 90 | 7.34 ± 0.21 | 7.64 ± 0.15 | 7.75 ± 0.13 |
| GαZ | 0 | 8.56 ± 0.11 | 8.98 ± 0.10 | 9.12 ± 0.10 |
|  | 90 | 7.48 ± 0.12 | 7.83 ± 0.14 | 7.97 ± 0.16 |

**Table S5:** CryoEM data collection, model refinement and validation of µOR- κ_ICL3_/naloxone/Nb6/10105-368-30-605 complex.

|  |  |
| --- | --- |
| **Data Collection** |  |
| Voltage (kV) | 300 |
| Magnification | 165,000x |
| Total electron dose ($\text{e}^-/\text{\AA}^2$) | 60 |
| Defocus range ($\mu\text{m}$) | -0.6 to -1.4 |
| Calibrated pixel size ($\text{\AA}$) | 0.741 |
| Micrographs collected | 10,007 |
| **Data Processing** |  |
| Extracted particles | 4,254,557 |
| Particles used for final reconstruction | 128,613 |
| Final map resolution ($\text{\AA}$, 0.143 FSC) | 3.26 |
| Map resolution range ($\text{\AA}$) | 2.8 – 3.6 |
| Map sharpening B factor ($\text{\AA}^2$) | 159.9 |
| **Model Content** |  |
| Initial models used (PDB code) | 7UL4 (μOR- $\kappa_{\text{ICL3}}$/Nb6/alvimopan) |
| Total number of atoms | 2546 |
| No. of protein residues | 339 |
| No. of ligands | 2 |
| **Model Validation** |  |
| CC map vs. model (%) | 0.73 |
| RMSD |  |
| Bond lengths ($\text{\AA}$) / Bond angles ($^\circ$) | 0.008/1.053 |
| Ramachandran plot statistics |  |
| Favored (%) | 93.60 |
| Allowed (%) | 6.40 |
| Outliers (%) | 0.00 |
| Rotamer outliers (%) | 0.00 |
| C-beta deviations | 0.00 |
| Clash score | 9.39 |

**Table S6:** 368 subtype selectivity by TRUPATH assay.

| Receptor/ligand | pEC<sub>50</sub> ± SEM (BRET-Gi1 protein activation), M |  |  |  |
| --- | --- | --- | --- | --- |
|  | 0 μM **368** | 3 μM **368** | 10 μM **368** | 30 μM **368** |
| μOR/DAMGO | 8.19 ± 0.06 | 8.11 ± 0.08 | 7.92 ± 0.09 | 7.62 ± 0.12 |
| δOR/DPDPE | 9.37 ± 0.05 | 9.24 ± 0.05 | 9.16 ± 0.06 | 8.91 ± 0.07 |
| κOR/U50488 | 8.77 ± 0.08 | 8.80 ± 0.08 | 8.68 ± 0.09 | 8.63 ± 0.11 |

**Table S7:**
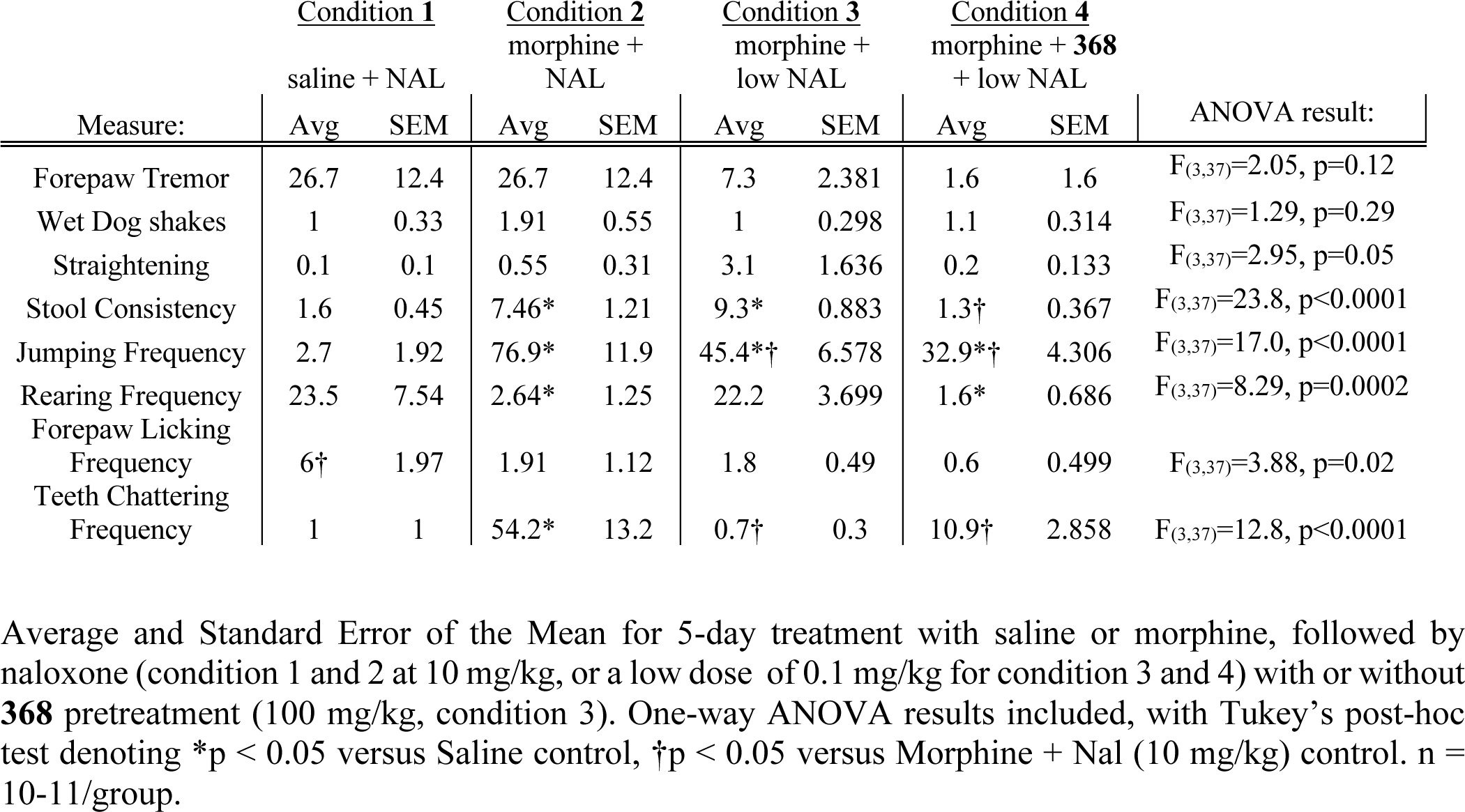
Summary of behavioral endpoints of naloxone-precipitated withdrawal in saline-treated or morphine-dependent mice following administration of either vehicle or 368 and naloxone (NAL)

